# Integration of the *Salmonella* Typhimurium methylome and transcriptome reveals DNA methylation and transcriptional regulation are largely decoupled under virulence-related conditions

**DOI:** 10.1101/2021.11.11.468322

**Authors:** Jeffrey S. Bourgeois, Caroline E. Anderson, Liuyang Wang, Jennifer L. Modliszewski, Wei Chen, Benjamin H. Schott, Nicolas Devos, Dennis C. Ko

**Affiliations:** Department of Molecular Genetics and Microbiology, School of Medicine, Duke University, Durham, NC, 27710, USA; University Program in Genetics and Genomics, Duke University, Durham, NC, 27710, USA; Center for Genomics and Computational Biology, Duke University, Durham, NC, 27710, USA; Division of Infectious Diseases, Department of Medicine, School of Medicine, Duke University, Durham, NC, 27710, USA

**Author notes:** To whom correspondence should be addressed: Dennis C. Ko, 0048B CARL Building Box 3053, 213 Research Drive, Durham, NC 27710. @denniskoHiHOST.

## Abstract

Despite being in a golden age of bacterial epigenomics, little work has systematically examined the plasticity and functional impacts of the bacterial DNA methylome. Here, we leveraged SMRT sequencing to examine the m^6^A DNA methylome of two *Salmonella enterica* ser. Typhimurium strains: 14028s and a Δ*metJ* mutant with derepressed methionine metabolism, grown in Luria Broth or a media that simulates the intracellular environment. We find that the methylome is remarkably static—over 95% of adenosine bases retain their methylation status across conditions. Integration of methylation with transcriptomic data revealed limited correlation between changes in methylation and gene expression. Further, examining the transcriptome in Δ*yhdJ* bacteria, lacking the m^6^A methylase with the most dynamic methylation pattern in our dataset, revealed little evidence of YhdJ-mediated gene regulation. Curiously, despite G(m^6^A)TC motifs being particularly resistant to change across conditions, incorporating *dam* mutants into our analyses revealed two examples where changes in methylation and transcription may be linked across conditions. This includes the novel finding that the Δ*metJ* motility defect may be partially driven by hypermethylation of the chemotaxis gene *tsr*. Together, these data redefine the *S*. Typhimurium epigenome as a highly stable system that has rare, but important, roles in transcriptional regulation. Incorporating these lessons into future studies will be critical as we progress through the epigenomic era.

**Importance:** While recent breakthroughs have enabled intense study of bacterial DNA modifications, limitations in current work have potentiated a surprisingly untested narrative that DNA methylation is a common mechanism of the bacterial response to environmental conditions. Essentially, whether epigenetic regulation of bacterial transcription is a common, generalizable phenomenon is a critical unanswered question that we address here. We find that most DNA methylation is static in *Salmonella enterica* serovar Typhimurium, even when the bacteria are grown under dramatically different conditions that cause broad changes in the transcriptome. Further, even when the methylation of individual bases change, these changes generally do not correlate with changes in gene expression. Finally, we demonstrate methods by which data can be stratified in order to identify coupled changes in methylation and gene expression.

## Introduction

Until recently, systematically understanding how the bacterial DNA methylome affects physiology has been an unachievable task. Unlike eukaryotes where m^**5**^C DNA methylation is highly abundant and can be detected with bisulfate sequencing (1), bacterial genomes primarily house m^6^A methylation which has historically been difficult to detect. Despite this technological hurdle, many studies over the last several decades have successfully uncovered roles for DNA methylation both in the contexts of restriction-modification systems (reviewed (2)), as well as for “orphan” methylases (particularly the Dam methylase) in DNA repair (3-9), DNA/bacterial replication and viability (10-19), *agn43* phase variation (20), LPS modifications (21-25), phage defense (21,26,27), mating (28,29), fimbriae formation (30,31), antibiotic resistance (32), hypoxia survival (33), motility (17,23,31,34), and other virulence related processes (8-10,16-19,23,31,34-39). While orphan methylases were originally thought to regulate bacterial physiology while restriction-modification systems targeted foreign DNA, recent work on “phasevarions” have found restriction-modification systems can indeed have dramatic impacts on the genome (reviewed (40)). A more complete history of associations between methylases and phenotypes can be found in recent reviews (41-43).

While early epigenome studies are valuable for the insights they provide, they depended on low-throughput and relatively blunt approaches (*e*.*g*., restriction enzyme digests paired with southern blotting to infer methylation). These approaches could not be leveraged to address if and how genome-wide changes in DNA methylation associate with changes to cellular processes. However, the discovery that sequencing data from the Pac-Bio SMRT-sequencing (44) and Oxford Nanopore sequencing (45,46) systems can be repurposed to detect m^6^A has heralded a golden age of bacterial DNA methylomics. These technological breakthroughs were rapidly applied to cataloging bacterial methylomes, many of which have been deposited in publicly available databases such as REBASE (47). However, we have only recently seen the power of these third-generation sequencing technologies applied to connect DNA methylation to cellular phenotypes. For instance, a recent paper utilized SMRT-seq to identify specific changes in G(m^6^A)TC patterns within the *opvAB* promoter that are highly present in the population following phage insult (48), building on previous phenotypic observations (21). Other groups have leveraged comparative epigenomics to examine methylation patterns across isolates and identify potentially important trends in methylation (35,49,50). Thus, there is immense potential for SMRT-seq to identify how methylation correlates with impactful biology.

Despite these advances we note that few studies have leveraged SMRT-seq to understand how methylation itself changes under different environmental pressures. Instead, the studies listed above typically examine methylomes under a single condition (typically late stationary-phase growth) to infer where methylation *can* happen, with notable exception (11,32,51-53). While informative, these approaches may not represent the methylation status of bacteria at growth phases typically studied in bacteriology, and thus may have limited ability to integrate into the broader microbiological literature and with transcriptomic datasets. A related shortcoming of many methylation studies is that methylation sites in promoters are often reported as evidence of methylation-mediated regulation, without determining whether methylation is dynamic under relevant environmental changes or testing whether disrupting methylation at those sites impacts transcription. Curiously, while a number of classical approaches have identified examples of methylation at specific sites regulating gene expression (*e*.*g. pap* (30,54,55), *opvAB* (21), *agn43* (56,57), *gtr* (22), the *std* operons (58), *dnaA* (12), *traJ* (29), and *sci1* (36)), we are unaware of any methylation site originally identified using genome-wide approaches that has been confirmed to impact gene expression. This disconnect between the technological advances in methylomics and the relatively modest conceptual advances in the field make it clear that the use of third generation sequencing technologies to interrogate the DNA methylome is still in its infancy and that the important question of the generalizability of epigenetic regulation of transcription in bacteria should be addressed.

In this paper, we perform a series of SMRT-seq and RNA-seq experiments to understand the role of DNA methylation in regulating *Salmonella enterica* serovar Typhimurium (*S*. Typhimurium) gene expression under environmental conditions critical for *Salmonella* virulence. Specifically, we studied conditions that activate the motility and *Salmonella* Pathogenicity Island-1 (SPI-1) pathways (growth in LB until late-log phase) and conditions that activate the *Salmonella* Pathogenicity Island-2 (SPI-2) pathways (growth in LPM media (59) until late-log phase). As methionine metabolism is intimately connected to methylation, we also examined the changes in methylation associated with derepressed methionine metabolism using a Δ*metJ* mutant. In general, we find that DNA methylation is mostly stable across conditions and is broadly decoupled from gene expression changes. This work redefines our understanding of the *S*. Typhimurium epigenome, provides multiple epigenomic datasets that can be incorporated into future work to identify rare instances where methylation changes are coupled with transcription, and a basic blueprint for carrying out future methylomic studies with high reproducibility.

## Materials and Methods

### Bacterial cell culture

All *Salmonella* strains are derived from *S*. Typhimurium NCTC 12023 (ATCC 14028s) and are included in **Supplemental Table 1**. All plasmids are included in **Supplemental Table 2**. Chromosomal knockouts were generated by lambda-red recombineering (60). Site-directed mutagenesis of the chromosome was performed using a modified version of lambda-red recombineering, as previously described (61). Complementation plasmids were generated by cut and paste cloning using the pWSK129 plasmid (62). For all experiments, bacteria were maintained on LB (BD, Miller formulation) agar plates, grown in LB media overnight at 37°C at 250 RPM, and subcultured the following morning prior to experiments. The SPI-2 inducing media is the low phosphate and magnesium (LPM) media from Coombes *et al*. (59) and contains 5 mM KCl, 7.5mM (NH_4_)_2_SO_4_, 0.5mM K_2_SO_4_, 38mM glycerol (0.3% volume/volume), 0.1% casein hydrolysate, 8μM MgCl_2_, 337μM K_2_HPO_4_ (pH 5.8), 80mM MES (pH 5.8), with the final solution pH equal to 5.8. Propagation of temperature sensitive plasmids occurred at 30°C and were cured at 42°C. Ampicillin was added to media at 100 μg/mL, kanamycin at 50 μg/mL, and apramycin at 100 μg/mL.

### Mammalian cell culture

THP-1 monocytes from the Duke Cell Culture Facility were cultured at 37°C in 5% CO2 in RPMI 1650 media (Invitrogen) supplemented with 10% heat-inactivated FBS, 2μM glutamine, 100 U/mL penicillin-G, and 100 mg/mL streptomycin. Cells used for *Salmonella* gentamicin protection assays were grown in antibiotic free media one hour prior to infection.

### Sample preparation for SMRT-Seq

*S*. Typhimurium were grown overnight, washed once, and subcultured 1:33 in LB for 2 hours and 45 minutes to induce SPI-1 expression, or 1:50 in SPI-2 inducing media for 4 hours in order to induce SPI-2 expression. 2×10^9^ bacteria were pelleted and DNA was extracted using a DNeasy Blood and Tissue kit (Qiagen). The optional RNase step in the protocol was performed to remove contaminating RNA, per manufacturer instructions. DNA was stored at 4°C until library preparation. Multiplexed SMRTbell libraries for sequencing on the PacBio Sequel system were prepared from 1 μg of each microbial gDNA sample. Shearing of gDNA was performed using g-TUBE and centrifugation at 2029 x g for 2 minutes to achieve a target mode size of 10 kb - 15 kb. SMRTbell libraries were then prepared using the SMRTbell Express Template Prep Kit 2.0. Two pools of 8 indexed libraries were prepared. Each pool was then sequenced on a PacBio Sequel SMRTcell using sequencing chemistry 3.0 and 10 hour movie length.

### Sample preparation for RNA-Seq

*S*. Typhimurium were grown overnight, washed once, and subcultured 1:33 in LB for 2 hours and 45 minutes, or 1:50 in SPI-2 inducing media for 4 hours. 2×10^9^ bacteria were pelleted at 5,000 x g for 5 minutes, and resuspended in RNAprotect Bacteria Reagent (Qiagen) in order to stabilize transcripts. After 5 minutes, bacteria were re-pelleted, and resuspended in 200 μL of TE Buffer containing lysozyme (15 mg/mL) and 20μL of Proteinase K. Bacteria were vortexed every two minutes for 15 minutes. 700 μL of β-mercaptoethanol-containing RLT buffer was added. After vortexing, 500 μL of 96% ethanol was added, and the solution was mixed and applied to a RNeasy extraction column (Qiagen). The remainder of the RNeasy protocol was followed per manufacturer instructions. After RNA isolation, 3-6 μg of RNA was treated with Turbo DNase (Thermo-Fisher) per manufacturer instructions, with the exception that two successive 30-minute DNase treatments were performed. To remove DNase after treatment, the solution was mixed with 350 μL of β-mercaptoethanol-containing RLT buffer, and then 700 μL of 96% ethanol was added. The mixture was then added to a RNeasy MinElute column (Qiagen) and RNA was reisolated according to manufacturer instructions.

RNA samples QC was performed with an Agilent Fragment Analyzer and a Qubit assay on the PerkinElmer Victor X2. Illumina TruSeq Stranded total RNA-Seq Kit combined with Ribo-Zero rRNA removal kit (bacteria) was used to prepare total RNA-seq libraries. Total RNA was first depleted of the rRNA using biotinylated probes that selectively bind to rRNA molecules. The rRNA depleted RNA was then reverse transcribed. During the 2nd strand synthesis, the cDNA:RNA hybrid is converted into to double-stranded cDNA (dscDNA) and dUTP incorporated into the 2nd cDNA strand, effectively marking the second strand. Illumina sequencing adapters were then ligated to the dscDNA fragments and amplified to produce the final RNA-seq library. The strand marked with dUTP is not amplified, allowing strand-specificity sequencing. Libraries were indexed using a dual indexing approach allowing for multiple libraries to be pooled and sequenced on the same sequencing flow cell of an Illumina MiSeq sequencing platform. Before pooling and sequencing, fragment length distribution and library quality was first assessed on a Fragment Analyzer using the High Sensitivity DNA Kit (Agilent Technologies). All libraries were then pooled in equimolar ratio and sequenced. Sequencing was done at 50 bp single-end reads. Once generated, sequence data was demultiplexed and Fastq files generated using Bcl2Fastq conversion software from Illumina.

### SMRT-seq mapping and m^6^A analysis

m^6^A methylation calls were performed using the pbsmrtpipe base modification and motif detection pipeline (Smrtlink v7.0.1.66975) with *Salmonella enterica* serovar Typhimurium strain 14028s (ASM2216v1) as the reference genome. For sites at or above 50x coverage, sites with a phred-based quality score greater than 40 were marked as “1”, for strong evidence of methylation; sites with >=50x coverage but below QV40 were marked as “0”, for unlikely to be methylated. For sites below 50x coverage, methylation status was not estimated. Assigning methylated bases to motif(s) was performed by comparing the context of the base to known or identified motifs using Microsoft Excel. Motif enrichment was calculated by dividing the frequency of the motif in a given subset (*e*.*g*. frequency of the motif in bases only methylated in bacteria grown in LB) and dividing by the frequency of the motif in condition tested (*e*.*g*. frequency of the motif among all methylated bases in bacteria grown in LB). Additional methyl-bases were detected on the pWSK29 plasmid harbored in these strains; however, we did not include these bases in our analyses as this plasmid is not involved in the natural lifestyle of *S*. Typhimurium.

### RNA-seq analysis and integration with methylomics

RNA-seq data was processed using the TrimGalore toolkit (http://www.bioinformatics.babraham.ac.uk/projects/trim_galore) which employs Cutadapt (63) to trim low quality bases and Illumina sequencing adapters from the 3’ end of the reads. Only reads that were 20 nt or longer after trimming were kept for further analysis. Reads were mapped to the ASM2216v1 version of the *Salmonella enterica* strain 14028S genome and transcriptome (64) using the STAR RNA-seq alignment tool (65). Reads were kept for subsequent analysis if they mapped to a single genomic location. Gene counts were compiled using the HTSeq tool (http://www-huber.embl.de/users/anders/HTSeq/). Only genes that had at least 10 reads in any given library were used in subsequent analysis. Normalization and differential expression was carried out using the DESeq2 (66) Bioconductor (67) package with the R statistical programming environment (https://www.R-project.org/).The false discovery rate was calculated to control for multiple hypothesis testing.

Integration of methylomics and RNA-seq analysis occurred in three steps. First, a list of genes present in both analyses was generated. Second, rates of differential expression among (a) the entire list of genes present in both analyses and (b) differentially methylated genes were called as genes containing 1+ base that was methylated in one condition but not another. Third, expected (frequency of differential expression in the entire list of genes present in both analyses multiplied by the frequency of differential methylation multiplied by the total number of genes in the analysis) and observed differentially methylated and differentially expressed genes were compared. Fisher’s Exact Test was used to determine whether there were statistically significant associations between differential methylation and differential expression.

### Analysis of *yhdJ* across *Salmonella* genomes

In order to analyze conservation of *yhdJ* across the *Salmonella enterica* genomes, 9,078 genomes (1,000 Typhimurium, 1,000 Typhi, 1,000 Paratyphi A, 1,000 Paratyphi B, 999 Newport, 1,000 Dublin, 1,000 Enteritidis, 1,000 Agona, 1,000 Heidelberg, and 79 Derby genomes) were obtained from the EnteroBase repository (68,69). Serovars examined here were specifically chosen to test for conservation among a diverse group of *Salmonella*. The specific strains were randomly selected and represented a variety of sources (human, agricultural animal, avian, reptiles, environment, etc.) within serovars, when possible. After downloading the genomes, all genomes of a given serovar were concatenated into a single FASTA file and used for analysis with the BLAST+ command line software (70). The 14028s YhdJ protein sequence was used as query for the pBLASTn program. To determine conservation, the program produced BLAST scores for ‘n’ sequences, where n = the number of strains tested within each serovar. The BLAST scores were then plotted relative to the BLAST score obtained using the 14028s genome.

### GO-term analysis

All GO-terms were generated using the Gene Ontology Resource (http://geneontology.org/) (71,72). The PANTHER Overrepresentation Test was run using the *Salmonella* Typhimurium GO biological process reference, the test used Fisher’s Exact test, and the correction was based on a calculated false discovery rate. All calculations were run automatically though the web portal software. Any gene that was not present in the GO-term database was “unmapped” and excluded from the analysis.

### Growth curves

*S*. Typhimurium were grown overnight in LB, subcultured 1:50 into 5mL of either LB or SPI-2 inducing media, and grown at 37°C at 250RPM. OD600 measurements were taken every 30 minutes using a spectrophotometer (Pharmacia Biotech Novaspec II).

### Gentamicin protection assay

Invasion and replication were measured as previously described (73-75). Briefly, bacteria were grown overnight, subcultured 1:33 into 1mL of LB, and grown for 2 hours and 45 minutes or until all strains entered late-log phase growth (OD600=1.5-2.0) at 37°C with 250 RPM. For any experiment using Δ*dam* bacteria, all bacteria were grown an extra 30 minutes (3 hours and 15 minutes) so that the Δ*dam* and Δ*dam*Δ*metJ* mutants reached late-exponential phase growth. 100,000 THP-1 monocytes, in antibiotic free media, were then infected by *S*. Typhimurium (MOI 5). At one hour post infection, cells were treated with gentamicin (50μg/mL), and IPTG was added 2 hours post infection to induce bacterial GFP expression. At 3 hours and 15 minutes post infection, cells were read by a Guava Easycyte Plus flow cytometer (Millipore). At 22 hours and 45 minutes post infection, IPTG was added to remaining wells to induce GFP, and at 24 hours post infection, cells were quantified by flow cytometry. Percent host cell invasion was determined by quantifying the number of GFP+ cells 3 hours and 15 minutes post infection, and replication was assessed by determining the ratio of the median intensity of GFP positive cells at 24 hours post infection divided by the median of the GFP positive cells at 3 hours and 15 minutes post infection.

### Motility assays

All strains were cultured overnight in LB, subcultured 1:33 into LB, and grown for 2 hours and 45 minutes at 37°C with 250 RPM. A pipette tip was used to puncture and deliver 2μL of *S*. Typhimurium into the center of a 0.3% LB agar plate. Plates were incubated at 37°C for 6 hours before the halo diameter was quantified.

### Murine competitive index experiments

Mouse studies were approved by the Duke Institutional Animal Care and Use Committee and adhere to the *Guide for the Care and Use of Laboratory Animals* of the National Institutes of Health. All experiments were performed with age- and sex-matched C57BL/6J (7-14 weeks old) mice. Bacteria were grown overnight, subcultured 1:33, and grown for 2 hours and 45 minutes at 37°C with 250 RPM. The bacteria were then washed and resuspended in PBS. Inoculums were confirmed by plating for CFUs. For oral infections, mice were fasted for 12 hours before infection, and given 100 μL of a 10% sodium bicarbonate solution by oral gavage 30 minutes before infection. Mice then received a 1:1 mixture of two *S*. Typhimurium strains containing either pWSK29 (AmpR) or pWSK129 (KanR) (62), totaling 10^8^ CFU in 100μL, by oral gavage. For intraperitoneal (IP) infections, mice were injected with a 1:1 mixture of two *S*. Typhimurium strains, totaling 10^3^ CFU in 100uL, into the intraperitoneal space. For both models, tissues were harvested four days post infection, homogenized, and plated on LB agar containing either ampicillin or kanamycin. Competitive index was calculated as (# Strain A CFUs in tissue/# Strain B CFUs in tissue)/ (# Strain A CFUs in inoculum/# Strain B CFUs in inoculum). Statistics were calculated by log transforming this ratio from each mouse and comparing to an expected value of 0 using a one-sample t-test.

### RT-qPCR

RNA was harvested as described above and used to create cDNA using the iScript cDNA synthesis kit (Bio-Rad Laboratories). qPCR was performed using the iTaq Universal SYBR Green Supermix (Bio-Rad Laboratories). 10 μL reactions contained 5 μL of the supermix, a final concentration of 500 nM of each primer, and 2 μL of cDNA. Reactions were run on a QuantStudio 3 thermo cycler. The cycling conditions were as follows: 95°C for 30 seconds, 40 cycles of 95 degrees for 15 seconds and 60°C for 60 seconds, and 60°C for 60 seconds. A melt curve was performed in order to verify single PCR products. The comparative threshold cycle (C_T_) method was used to quantify transcripts, with the ribosomal *rrs* gene serving as the endogenous control. Fold change represents 2^-ΔΔCT^. Oligonucleotides are listed in **Supplemental Table 3**.

### Western blotting

*flhC* was tagged with the 3xFLAG tag using recombineering as previously described (76). *S*. Typhimurium were grown overnight in LB, subcultured 1:33 in LB at 37°C with 250 RPM until late log phase (OD600-1.5-2.0), and pelleted by centrifugation at 6,000 x g for 5 minutes. Pellets were resuspended in 2x laemmli buffer (Bio-Rad) with 5% 2-Mercaptoethanol, boiled for 10 minutes, and lysates were run on Mini-PROTEAN TGX Stain-Free gels (Bio-Rad). After electrophoresis the gels’ total protein dye was activated by a 5-minute UV exposure. Following transfer onto Immun-Blot low-fluorescence PVDF membrane (Bio-Rad) using a Hoefer TE77X, blots were probed using an anti-FLAG M2 antibody (Sigma F3165). A florescent secondary antibody (LI-COR IRDye) was used to detect bands on a LI-COR Odyssey Classic. Band intensity was quantified using LI-COR Odyssey Imaging System Software v3.0. Total protein was detected by 30 seconds of UV exposure, and quantified using Fiji (77). The graphed relative signal is: (FLAG band intensity/Total Protein) divided by (FLAG band intensity in wild-type *flhC:FLAG3x* bacteria/Total Protein in wild-type *flhC:FLAG3x* bacteria).

### Statistical analyses

Statistics were performed in Graphpad Prism 9 or Microsoft Excel, except where otherwise noted. Where noted, inter-experimental noise was removed from gentamicin protection assays and motility assays prior to data visualization or statistical analysis by standardizing data to the grand mean by multiplying values within an experiment by a constant (average of all experiments divided by average of specific experiment). All statistical tests corresponding to reported p-values are described in the appropriate figure legends.

## Results

### A genome-wide screen to understand how growth conditions and methionine metabolism impact m^6^A DNA methylation

While previous work on bacterial DNA methylation has largely focused either on global DNA methylation patterns under a single condition or on how methylation of a single motif changes under different conditions, we sought to examine how the entire *S*. Typhimurium m^6^A DNA methylome changes under four biologically relevant conditions **(Figure 1A)**. We examined aerobic growth in LB media to late exponential phase (OD600 ∼1.5-2.0), which induces expression of flagellar genes and the genes in the *Salmonella* Pathogenicity Island-1 (SPI-1)—including the type III secretion system used during host cell invasion (78). The second condition cultured bacteria in a minimal media used to induce expression of genes in the *Salmonella* Pathogenicity Island-2 (SPI-2) (59)—which include the type III secretion system turned on in the host cell to promote *Salmonella* vacuolar survival (79,80). The third and fourth conditions repeated growth in these media but used a methionine metabolism mutant *S*. Typhimurium strain, Δ*metJ*. The MetJ protein represses expression of methionine metabolism genes (81). Thus, Δ*metJ* bacteria have deregulated methionine metabolism, and accumulation of methionine and related metabolites, including metabolites directly related to methylation processes such as the universal methyl-donor S-adenosyl-methionine (SAM) (82), and the methyltransferase-inhibiting metabolites methylthioadenosine (83-85) and S-adenosyl-homocysteine (86-90). Of note, we have not only previously confirmed increased abundances of both SAM and MTA in Δ*metJ*, but also demonstrated that the Δ*metJ* mutant has attenuated SPI-1 secretion, motility, and virulence (91) and had previously hypothesized that these effects could be mediated through aberrant methylation.

**Figure 1:**
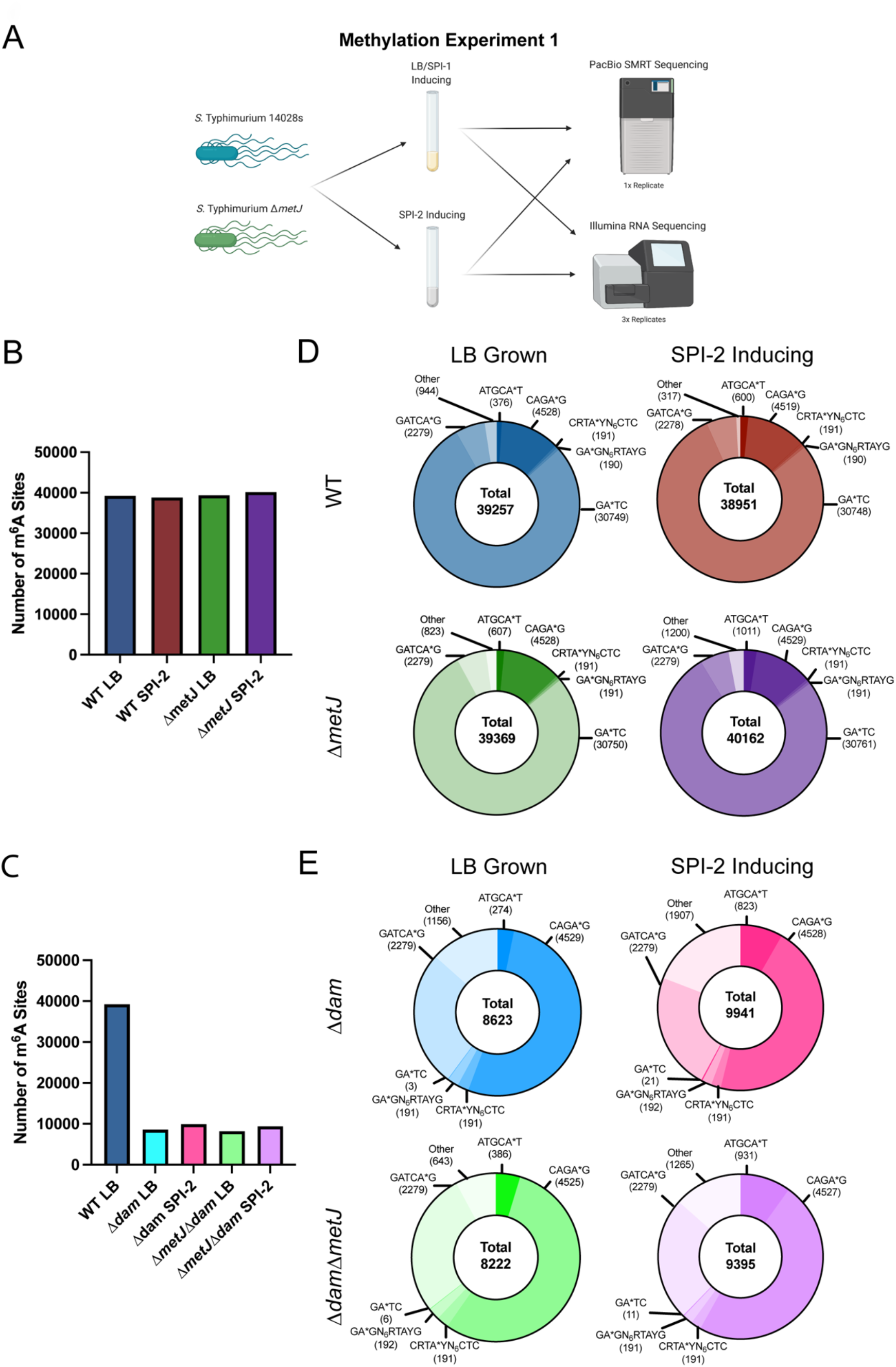
Genome-wide analysis of m^6^A DNA methylation under varying conditions. (A) Schematic of Methylation Experiment 1. Wild-type *S*. Typhimurium (Strain 14028s) and an isogenic Δ*metJ* strain were cultured in LB or *Salmonella* Pathogenicity Island-2 (SPI-2)-inducing media and DNA was collected for SMRT-sequencing. Bacteria grown under identical conditions were harvested for RNA-sequencing. (B, C) Total number of m^6^A bases observed across conditions does not dramatically change in wild-type and Δ*metJ* bacteria (B), but does change dramatically in Δ*dam* bacteria (C). (D) Analysis of motifs methylated reveals only the total number of ATGCA*T and “other” sites (sites that do not map to one of the six motifs) changes dramatically across conditions. (E) Δ*dam* results in ablation of GATC methylation. For Panels B through E, bases were only included in the analysis if the base could confidently be called methylated or unmethylated across the eight conditions.

To analyze the DNA methylome, we performed a PacBio SMRT-sequencing experiment (hereon called “Methylation Experiment 1”) in biological singlet (as has been common in the field and as we comment on below) to identify whether any changes in methylation could be observed. In this experiment, we also included Δ*dam* and Δ*dam*Δ*metJ* mutants, which lack G(m^6^A)TC (henceforth the * symbol will denote the adenosine that is m^6^A modified; GA*TC) methylation grown under SPI-1 and SPI-2 conditions. This allowed us to confirm that our pipeline could adequately detect changes in methylation. These eight conditions were split across two PacBio SMRT Cells. Thus, these conditions enabled comparison of the *S*. Typhimurium DNA methylome under the two conditions most critical for *Salmonella* virulence, under perturbation of methionine metabolism, and with a control condition ablating the primary DNA methyltransferase.

In total this experiment defined the methylation status of 61,704 adenosine bases **(GEO: GSE185578)**, however, methylation status of some bases under certain conditions could not be determined as coverage was below 50X. Thus, we restricted our analysis to 51,177 bases in which the methylation status could be adequately determined for all conditions tested. These bases span both the *S*. Typhimurium genome and virulence plasmid.

To compare methylation across conditions, we called methylation in two ways. First, we assigned each base a “percent methylated” value, which considered the percent of reads for each base that were counted as methylated compared to the total number of reads **(Supplemental File 1)**. We also examined the data as a binary in which we considered bases either methylated (if any methylation was detected) or unmethylated **(Supplemental File 2)**. Using this binary analysis, we observed that there were similar, but subtly different, amounts of m^6^A methylation across wild-type and Δ*metJ* bacteria grown in LB and SPI-2 conditions (WT LB: 39,240 bases; WT SPI-2: 38,827 bases; Δ*metJ* SPI-1: 39,352 bases; Δ*metJ* SPI-2: 40,145 bases) **(Figure 1B)**, but that, as expected, Δ*dam* reduced total methylation substantially **(Figure 1C)**.

### ATGCAT motifs are frequently differentially methylated across conditions

We next examined how these bases were distributed across different methylation motifs. This analysis detected methylation at motifs that had been previously detected in *S*. Typhimurium (92), though we were able to detect an additional motif, CRTA*YN_6_CTC, which appears to be the reverse complement of GA*GN_6_RTAYG. Notably, two motifs (CAGA*G and GA*GN_6_RTAYG) cannot always be distinguished, and so we included bases that matched to both motifs in counts for each. Bases that did not map to any known motif were listed as “Other.” As with the total amount of m^6^A methylation, we found that our four main conditions had subtle differences in the total numbers of most motifs **(Figure 1D)**.

The most notable change in motif abundance occurred at the ATGCA*T motif, which is methylated by the YhdJ methylase (93). We observed more ATGCA*T methylation in bacteria grown under SPI-2-inducing conditions (p<0.00001, Chi-Square Test) or in Δ*metJ* bacteria (p<0.00001, Chi-Square Test), with the highest ATGCA*T methylation present in Δ*metJ* bacteria grown under SPI-2-inducing conditions. We also observed variation in the number of bases that mapped to the “Other” category. In contrast, we observed very little change in the total amount of GA*TC methylation (methylated by Dam) across these physiologically relevant conditions, though deletion of *dam* resulted in almost complete ablation of methylation at the GATC motif **(Figure 1E)**.

To examine the methylation changes under LB vs. SPI-2 inducing conditions with wild-type and Δ*metJ S*. Typhimurium, we compared binary methylation at each individual base to identify differentially methylated bases (bases that were called methylated in one condition but not another). While each condition had a few hundred to over a thousand bases that were not methylated in their opposing group, the vast majority of bases in this study (>38,000) were shared across these comparisons **(Figure 2A, 2B, 2C; Venn Diagrams)**. This demonstrates that while the methylome is slightly responsive to the environment and methionine metabolism, it remains largely static across strikingly different conditions.

**Figure 2:**
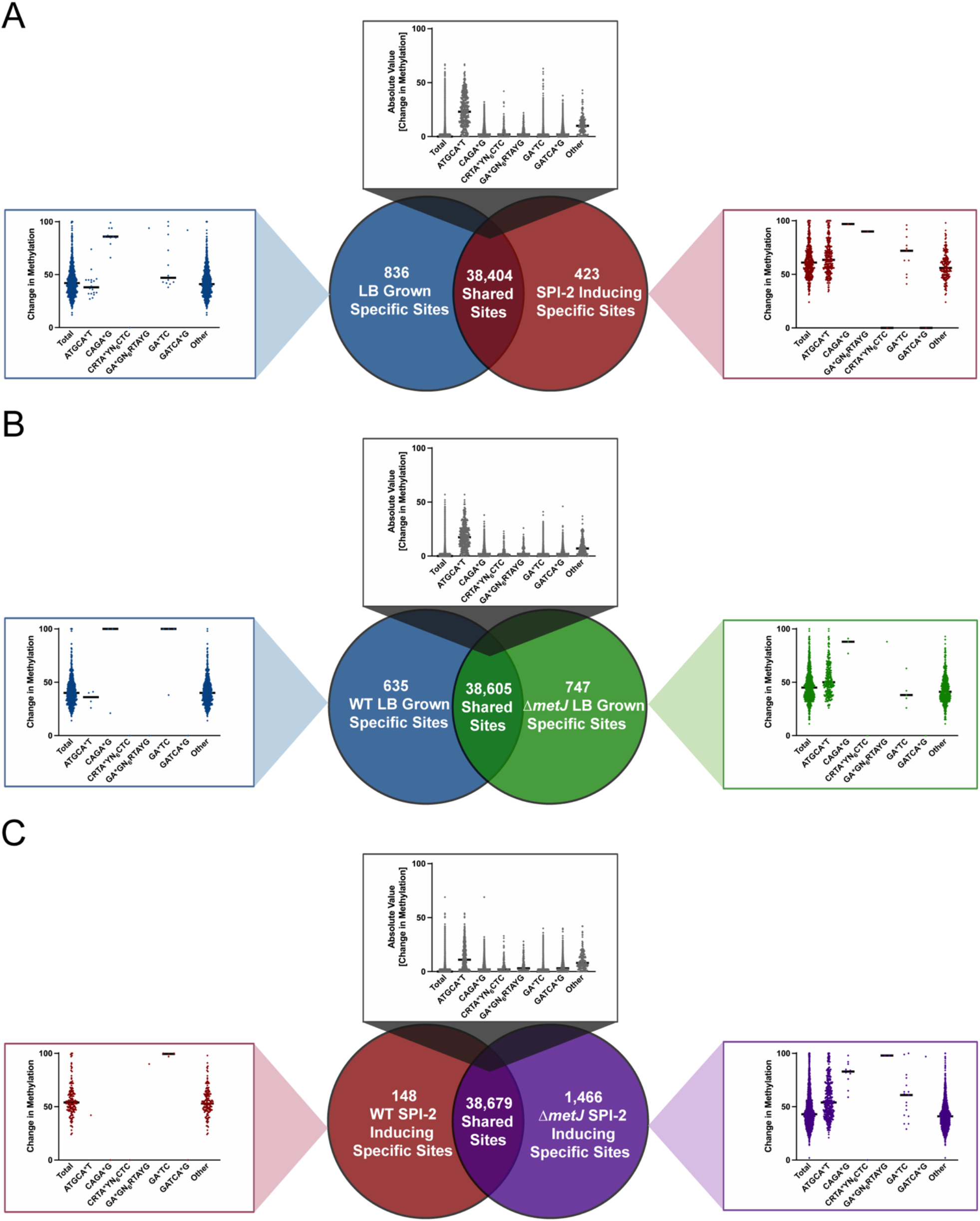
Integration of binary and quantitative analyses to understand differential methylation in *S*. Typhimurium. (A-C) Quantification of shared and unique methylated sites between wild-type *S*. Typhimurium grown in LB and SPI-2 inducing media (A), WT and Δ*metJ* bacteria grown in LB (B), and WT and Δ*metJ* bacteria grown in SPI-2 inducing media (C). Venn diagrams are based on binary measures of differential methylation. Sites identified by the binary analysis were examined in our quantitative dataset in order to identify changes in the percent methylation. In the graphs “Total” refers to all sites present in the relevant part of the Venn diagram, which were then broken down by motif. For motifs where no differentially methylated sites were present, a single dot is listed at 0%. For shared sites, the absolute value of the difference between bases are shown and thus the numbers are agnostic to whether methylation is higher in either condition. Bars mark the median. For all panels, only bases that could be confidently called methylated or unmethylated in the eight conditions in **Figure 1** were considered.

Having identified these differentially methylated bases by our binary analyses, we integrated our quantitative data. This is an important measurement as previous work has speculated that methylation impacts bistable gene expression (48,94), and thus changes in the percent of the population in which a given base is methylated could have implications on the percent of the population expressing a given gene. For each differentially methylated base, we asked what the total change in methylation was across the two conditions **(Figure 2A, 2B, 2C; Graphs)**. The total median shift in the percent methylation varied by condition, but fell between 43% and 53%, suggesting that most bases go from unmethylated in one condition, to about half the copies of the genome having methylation at that site in the other. Notably, most of the “shared” bases that demonstrated methylation under both conditions demonstrated no quantitative change (median = 0%). However, again the exception was ATGCA*T, where the median shift among shared bases remained relatively high (11-23% depending on condition).

To test for enrichment of motifs among differentially methylated bases, we compared the frequency of each of the six motifs tested above in the differentially methylated sites against the frequency observed in the entire condition. This analysis revealed that among the uniquely SPI-2-induced methylated bases, we observed 37 times more differentially methylated ATGCA*T sites than expected. Similarly, sites methylated in Δ*metJ S*. Typhimurium, but not wild-type bacteria, grown under both LB and SPI-2-inducing conditions are also dramatically enriched for YhdJ-mediated methylation (20-fold and 11-fold enrichment, accordingly) **(Supplemental Figure 1)**. Surprisingly, all other motifs were either present at similar or dramatically lower abundance among differentially methylated sites than expected by chance, though we did note significant enrichment of “other” motifs among differentially methylated bases (20 to 100-fold enrichment, depending on condition).

### A replication experiment demonstrates that SMRT-seq is highly reproducible and confirms differential methylation is predominantly driven by YhdJ

To confirm our findings that DNA methylation was largely stable among our conditions with the exception of ATGCA*T sites, we repeated our SMRT-seq experiment with wild-type and Δ*metJ* bacteria grown in LB **(Supplemental Figure 2A;** Replication Methylation Experiment). Further, to confirm that the significant enrichment in ATGCA*T methylation in Δ*metJ* bacteria we observed above was due to YhdJ, we also sequenced Δ*yhdJ* and Δ*yhdJ*Δ*metJ S*. Typhimurium grown in LB. Of note, while we sequenced eight samples across two SMRT cells in Methylation Experiment 1, here we sequenced these four samples on two SMRT cells, significantly increasing our sequencing depth. The resulting dataset called the methylation status of 60,502 bases in at least one condition, and, strikingly, 60,501 of these bases were confidently called in all four conditions **(GEO: GSE185501**).

Analysis of the two methylomic datasets using our binary assessment revealed that ∼97.5% of bases replicated their methylation status across experiments, demonstrating that our results were highly reproducible **(Supplemental Figure 2B, Supplemental File 2)**. Importantly, we again observed that Δ*metJ* bacteria have increased ATGCA*T methylation and found 0 methylated ATGCA*T sites in the Δ*yhdJ* and Δ*yhdJ*Δ*metJ* mutants, confirming that YhdJ is the only ATGCA*T methylase active in both bacterial strains **(Supplemental Figure 2C)**. Of note, sites assigned to “other” motifs may include significantly more miscalled methyl bases, as only ∼80% (79.3% of wild-type and 80.1% of Δ*metJ*) “other” bases replicated their methylation status across experiments.

Analysis of the two datasets using the quantitative measurement **(Supplemental File 3)** revealed considerable replication across the datasets (wild-type R^2^=0.87 and Δ*metJ* R^2^=0.86) **(Supplemental Figure 2D)**. Considering these experiments as separate biological replicates and using an arbitrary cutoff of 10% average differential methylation, we identified 2,528 sites (out of 50,962 total sites; 4.96%) that were differentially methylated between wild-type and Δ*metJ* bacteria using this quantitative method (**Supplemental Figure 2E)**. 881 of these sites were more methylated in wild-type bacteria, and 1,647 were more methylated in Δ*metJ* bacteria.

Having assessed the reproducibility of SMRT-Seq for both categorical and quantitative measures of methylation, we used our binary measurement to generate a combined dataset containing bases which were (a) reliably detected in wild-type and Δ*metJ* bacteria grown in LB in both experiments, and (b) were identically called methylated or unmethylated in both experiments. Using this dataset (52,594 bases), we determined which differentially methylated bases repeated across the two studies. This number of bases is greater than the number of bases included in Methylation Experiment 1 analyses, as we no longer needed to exclude bases that did not reach sufficient coverage in either the SPI-2 inducing or Δ*dam* conditions. While our data demonstrated that the vast majority of bases were called identically (∼97.5%; **Supplemental Figure 2B)**, we found that a disproportional number of bases that were called differentially methylated in the pilot study failed to replicate in the replication study, and vice versa. In fact, while there were 1,382 bases called differentially methylated in the first experiment **(Figure 2B)**, and 2,544 bases called differentially methylated in the replicate study **(Supplemental Figure 2F)**, only 308 differentially methylated bases were identified in the combined dataset (**Supplemental Figure 2G)**. Importantly, the overlap between these two replicates is much greater than expected by chance (3.7-fold enrichment; p<0.0001, one-tailed binomial test), indicating that these biological replicates provide a high-confidence set of 308 differentially methylated sites, though some false positives likely remain. Our findings emphasize that while SMRT-seq calling of methylated bases is reliable, replication is especially important in examining bases that change between conditions. Of note, the combined dataset once again revealed enrichment of differentially methylated “other” sites. To understand these sites, we examined the 40 bases surrounding the 143 instances of “other” differential methylation in the Combined Dataset using the Multiple Em for Motif Elicitation (MEME) software (95). This identified a single significant motif (E-value=6.1×10^−16^), ACCWGG **(Supplemental Figure 3A)**. The same motif was identified among the 969 differentially methylated “other” sites between LB and SPI-2 grown bacteria (Methylation Experiment 1; E-value = 5.5×10^−206^; **Supplemental Figure 3B**). The ACCTGG motif has been reported on multiple *Salmonella* serovar entry pages on REBASE (47), however, curators note that this is almost certainly a miscall for the m^5^C motif CCWGG—methylated by Dcm. Across the combined dataset, we found 33 instances of this motif (23% of all differentially methylated “other” sites). This leads us to hypothesize that this dynamic “other” category may be predominantly driven by changes in the flexible m^5^C methylome, which warrants further investigation using sequencing technologies better equipped to detect cytosine methylation.

### There is no association between the transcriptome and the genome-wide binary methylation analyses under the conditions tested here

Canonically, changes in DNA methylation can lead to changes in transcription by enabling differential binding of transcription factors to genomic elements (reviewed (41)). However, studies that describe this in bacteria typically either (a) focus on single loci (for example, (48)), or (b) only speculate on direct mechanisms of transcriptional control based on methylase knockout experiments (for example (35)). No study has directly examined whether differential methylation across the *S*. Typhimurium genome correlates with differential expression under biologically relevant conditions. We attempted to fill this gap in knowledge by performing RNA-seq on wild-type and Δ*metJ* bacteria grown in LB and SPI-2 inducing media **(GEO: GSE185072, Supplemental File 4)** and looking for correlations with our SMRT-seq datasets.

Prior to integrating our datasets, we confirmed our RNA-seq data matched previously observed trends. As expected, we identified many differentially expressed genes (2,639 DEGs at FDR < 0.5) between wild-type bacteria grown in LB vs. SPI-2-inducing conditions **(Figure 3A)**. These DEGs included a variety of expected genes, including higher expression of SPI-1genes (*e*.*g*., *sipB*, 322-fold induction) and flagellar genes (*e*.*g*., *fliD*, 138-fold induction) in LB and higher expression of SPI-2 genes (*e*.*g*., *ssaN, sscA;* 182- and 579-fold induction respectively) in SPI-2 inducing media. Further, these data cluster with previous transcriptomic analyses of gene expression in LB and SPI-2 inducing conditions (96) by PCA (53.5% of variation between the two studies explained by media, **Supplemental Figure 4**). Analysis of Δ*metJ* DEGs revealed a number of expected trends, specifically that during growth in LB and SPI-2, Δ*metJ* shows upregulation of methionine metabolism genes resulting from direct derepression of the metabolic pathway **(Figure 3B, 3C)** (81). Of note, we also observed reduced motility gene expression which we had previously reported (91), but were surprised that contrary to our prior work we observed a small reduction in *flhD* expression in Δ*metJ*. However, we confirmed this result by qPCR and western blotting **(Supplemental Figure 5A, 5B)**, and speculate that improved DNase treatment in this study likely explains this difference.

**Figure 3:**
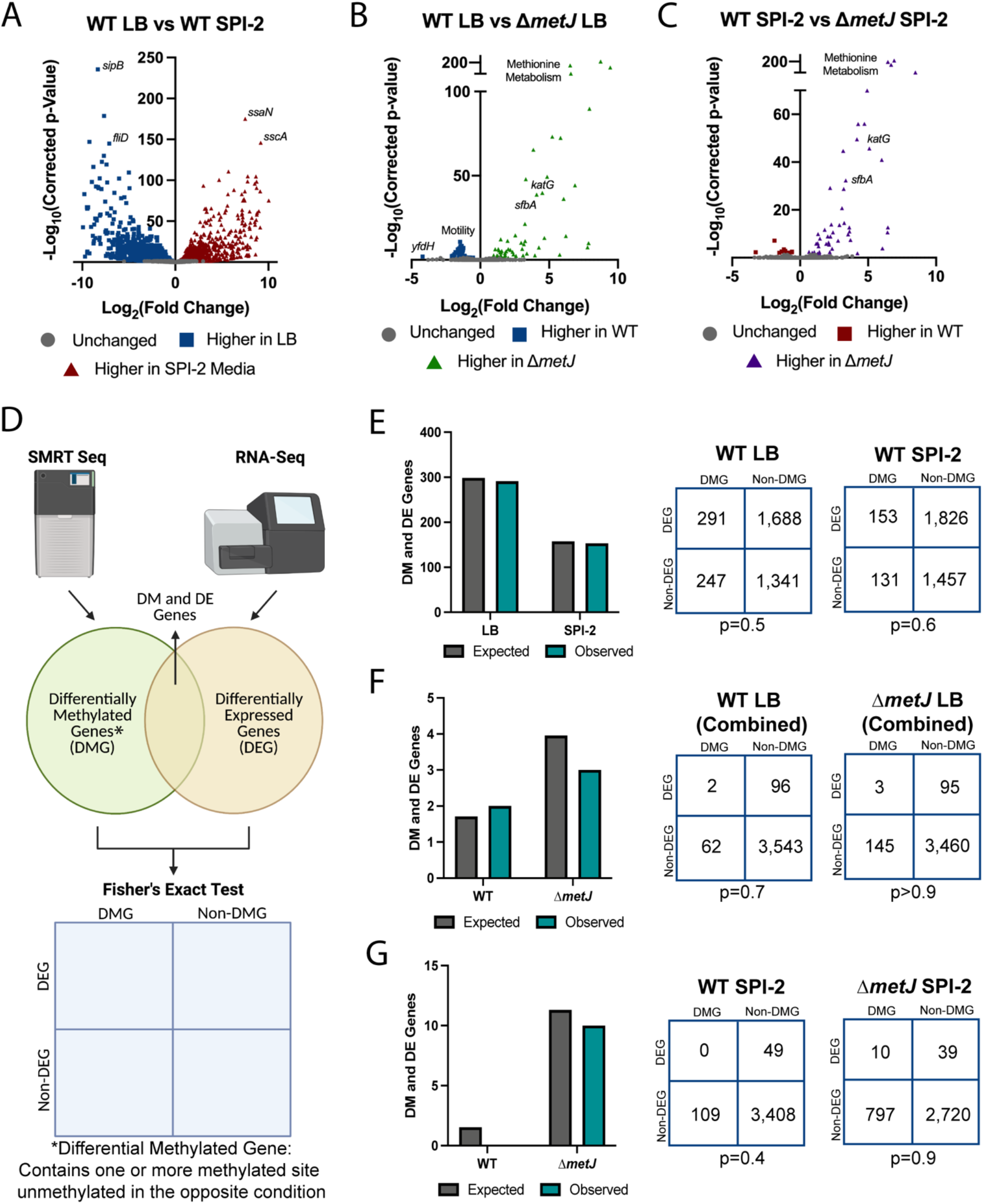
Differentially methylated genes by binary analysis are not enriched for transcriptomic changes. (A-C) RNA-seq reveals transcriptomic changes between LB grown and SPI-2 media grown wild-type bacteria (A), wild-type and Δ*metJ* bacteria grown in LB (B), and wild-type and Δ*metJ* bacteria grown in SPI-2 media (C). Corrected p-values generated by calculating the false discovery rate. (D) Schematic of RNA-seq and SMRT-seq integration. Each gene was determined to be differentially methylated (Differentially Methylated Gene, DMG) in our binary analysis, differentially expressed (Differentially Expressed Gene, DEG (FDR<0.05)), differentially methylated and differentially expressed (DM and DE Gene), or neither differentially methylated nor expressed. Fisher’s Exact Test was then used to determine whether there was an association between methylation and gene expression. (E-G) Differential methylation is not associated with differential expression. Observed and expected numbers of differentially methylated and differentially expressed genes were not significantly different when comparing uniquely methylated genes in LB vs SPI-2 media (E), wild-type vs Δ*metJ* in LB (F), or wild-type vs *ΔmetJ* in SPI-2 media (G). Uniquely methylated genes are plotted in the condition under which they are methylated (*e*.*g*. for panel E, a gene that contains a base that is methylated in LB but not SPI-2 media would be plotted as part of “LB”), but are agnostic to the direction of effect for the expression data. Expected values are calculated by multiplying the frequency of differential methylation by the frequency of differential expression by the total number of genes in the analysis for each condition. Numbers used for Fisher’s Exact Test are shown on the right. Data for E and G used data from Methylation Experiment 1, F used the “Combined Dataset.” For F and G the gene *metJ* is removed from the analysis, as it is artificially called both differentially methylated and expressed due to its excision from the genome.

To integrate our differential expression data with our methylomics data, we considered genes that either (a) contained or (b) were the closest gene to one or more binary differentially methylated sites (*ie*., present at any level in one condition, absent in the other) to be “differentially methylated genes” (DMGs) **(Figure 3D)**. For each comparison, the status “differentially methylated” applied to the condition in which the methyl mark was present (*e*.*g*. when comparing LB grown vs SPI-2 grown bacteria, an LB-grown DMG contains a methyl mark that is absent in SPI-2 media grown bacteria). Using these criteria, we examined whether differentially methylated genes were more likely to be differentially expressed than predicted by chance. Strikingly, we did not observe enrichment of DEGs among our DMGs. The number of DEGs that were also DMGs under all comparisons was remarkably similar to the overlap of these categories expected by chance (Fisher’s p-value > 0.05 in all cases), suggesting the two phenomena are typically not associated at the genome-wide level across any of our comparisons with binary calling of DMGs (**Figure 3E-G)**. Most importantly, there was no evidence of enrichment comparing WT *S*. Typhimurium grown in LB (SPI-1 inducing) and SPI-2 conditions (**Figure 3E**), indicating that co-occurrence of differential methylation and differential gene expression is not observed more frequently than is expected by chance in switching between these two critical growth conditions in *Salmonella* pathogenesis. Notably, we also did not observe correlations if we adjusted the statistical thresholds for differential expression (**Supplemental Figure 6A-D**), or if we stratified our data by the differential expression direction of effect **(Supplemental Figure 6E, 6F)**, the genic location of the differential methylation **(Supplemental Figure 6G)**, or to specific motifs **(Supplemental Figure 6H, 6I)**.

### There is limited association between the transcriptome and the genome-wide quantitative methylation analyses under the conditions tested here

In addition to these binary definitions to differential methylation, we examined whether there was enrichment of DEGs among DMGs defined by a difference in ≥10% methylation across conditions **(Figure 4A)**. Most of our binary observations replicated in this analysis **(Figure 4B, 4C)**, but we noted an association between DEGs and DMGs in wild-type bacteria grown in SPI-2 inducing media compared to Δ*metJ* bacteria grown in the same conditions **(Figure 4D, Table 1)**. It is unclear why this condition is the exception to the general lack of association we observe, but it may suggest that methylases (particularly the *dam* methylase) and deregulated transcriptional machinery uniquely compete for access to these sites exclusively in minimal media. We also examined whether adjusting our thresholds for differential expression or limiting our search for differential to bases upstream of differentially expressed genes could reveal further correlations between expression and methylation, but no additional associations were found **(Supplemental Figure 7A-F)**. The association with sites in wild-type bacteria in SPI-2 media compared to Δ*metJ* was still present with the more stringent DEG cutoff **(Supplemental Figure 7C)** but was no longer present when analysis was restricted to upstream bases **(Supplemental Figure 7F)**. Finally, we leveraged the quantitative dataset to examine the relationship between DMGs and DEGs in which DMGs are defined by bases completely methylated in one condition (≥99%) and hypomethylated in the other (≤89%) (**Supplemental Figure 7G-I)**. This revealed no additional associations, but the association with wild-type bacteria in SPI-2 media replicated once again. We conclude that for the crucial switch between SPI-1 and SPI-2 virulence gene programs, there is no association between m^6^A DNA methylation and transcriptional regulation but that specific mutants may show an association.

**Figure 4:**
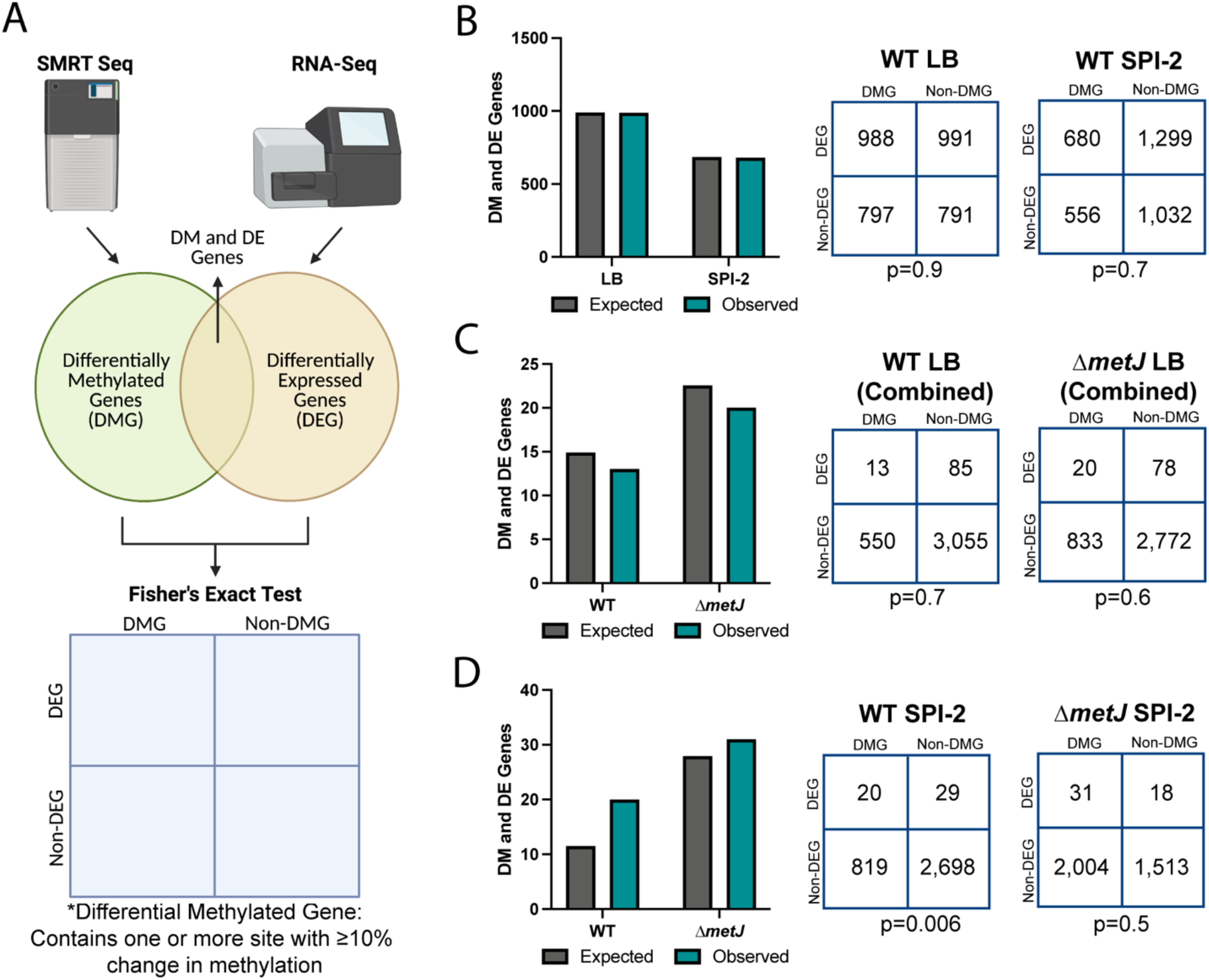
Quantitative analysis revealed an association between differential methylation and expression between wild-type and Δ*metJ* bacteria. (A) Schematic of RNA-seq and SMRT-seq integration. Each gene in our quantitative analysis was determined to be differentially methylated (Differentially Methylated Gene, DMG: difference ≥10% methylation across conditions), differentially expressed (Differentially Expressed Gene, DEG: FDR corrected p-value ≤0.05), differentially methylated and differentially expressed (DM and DE Gene), or neither differentially methylated nor expressed. Fisher’s Exact Test was then used to determine whether there was an association between methylation and gene expression. (B-D) Differential methylation is typically not associated with differential expression. Observed and expected numbers of differentially methylated and differentially expressed genes were not significantly different when comparing uniquely methylated genes in LB vs SPI-2 media (B), wild-type vs Δ*metJ* in LB (C), however, a significant enrichment of DEGs was observed in hypermethylated sites in wild-type bacteria grown in SPI-2 media relative to *ΔmetJ* (D). Hypermethylated genes are plotted in the condition under which they have increased methylation (*e*.*g*. for panel B, a gene that contains a base that is more methylated in LB would be plotted as part of “LB”), but are agnostic to the direction of effect for the expression data. Expected values are calculated by multiplying the frequency of differential methylation by the frequency of differential expression by the total number of genes in the analysis for each condition. Numbers used for Fisher’s Exact Test are shown on the right. Data for B and D used data from Methylation Experiment 1, C used the “Combined Dataset.” For C and D the gene *metJ* is removed from the analysis, as it is excised from the genome.

**Table 1:** Correlation of differential methylation and differential expression in wild-type *S*. Typhimurium grown in SPI-2 inducing media compared to Δ*metJ* bacteria.

| Genomic Location | WT (SPI-2) % Methylated | $\Delta metJ$ (SPI-2) % Methylated | $\Delta$ %Methylation (WT-$\Delta metJ$) | STM14_ID | Common Name | Relative Location | Motif | log<sub>2</sub>FoldChange ($\Delta metJ$_SPI-2 / WT_SPI-2) |
| --- | --- | --- | --- | --- | --- | --- | --- | --- |
| NC_016856.1__1217 | 99 | 88 | 11 | STM14_0002 | thrA | Genic | GATCA*G | -1.35 |
| NC_016856.1__277289 | 99 | 88 | 11 | STM14_0276 | ldcC | Genic | GA*TC | 0.89 |
| NC_016856.1__288405 | 98 | 87 | 11 | STM14_0289 | metN | Genic | GA*TC | 4.27 |
| NC_016856.1__572813 | 93 | 74 | 19 | STM14_0601 | sfbB | Genic | CAGA*G | 3.25 |
| NC_016856.1__573231 | 100 | 90 | 10 | STM14_0601 | sfbB | Genic | GA*TC | 3.25 |
| NC_016856.1__573265 | 100 | 81 | 19 | STM14_0601 | sfbB | Genic | GA*TC | 3.25 |
| NC_016856.1__573302 | 100 | 88 | 12 | STM14_0601 | sfbB | Genic | GA*TC | 3.25 |
| NC_016856.1__573803 | 100 | 90 | 10 | STM14_0602 | NA | Genic | GA*TC | 3.62 |
| NC_016856.1__861113 | 100 | 87 | 13 | STM14_0920 | bioB | Genic | GA*TC | -1.22 |
| NC_016856.1__861413 | 100 | 88 | 12 | STM14_0920 | bioB | Genic | GA*TC | -1.22 |
| NC_016856.1__1048140 | 98 | 88 | 10 | STM14_1130 | ompF | Genic | GA*TC | 1.32 |
| NC_016856.1__1423354 | 100 | 89 | 11 | STM14_1619 | thrS | Genic | GA*TC | -0.59 |
| NC_016856.1__1733156 | 76 | 40 | 36 | STM14_1975 | NA | Upstream | Other | 1.46 |
| NC_016856.1__1829331 | 99 | 86 | 13 | STM14_2086 | NA | Genic | GA*TC | 3.63 |
| NC_016856.1__1831286 | 100 | 71 | 29 | STM14_2087 | trpD | Genic | GA*TC | 3.71 |
| NC_016856.1__1834122 | 100 | 84 | 16 | STM14_2089 | trpB | Genic | CAGA*G | 2.13 |
| NC_016856.1__2340851 | 98 | 81 | 17 | STM14_2702 | NA | Genic | GA*TC | 2.79 |
| NC_016856.1__2746393 | 100 | 88 | 12 | STM14_3133 | NA | Genic | GATCA*G | 1.80 |
| NC_016856.1__3270917 | 100 | 88 | 12 | STM14_3733 | metK | Genic | GA*TC | 3.21 |
| NC_016856.1__3272719 | 100 | 88 | 12 | STM14_3735 | NA | Genic | GA*TC | -0.82 |
| NC_016856.1__3722412 | 100 | 90 | 10 | STM14_4260 | asd | Genic | CAGA*G | 2.03 |
| NC_016856.1__4334354 | 100 | 87 | 13 | STM14_4936 | katG | Genic | GA*TC | 5.10 |
| NC_016856.1__4417534 | 100 | 90 | 10 | STM14_5030 | aceA | Genic | GA*TC | 2.42 |
| NC_016856.1__4417670 | 100 | 78 | 22 | STM14_5030 | aceA | Genic | GA*TC | 2.42 |
| NC_016856.1__4423175 | 100 | 89 | 11 | STM14_5035 | metH | Genic | GA*TC | 3.09 |
| NC_016856.1__4424372 | 100 | 89 | 11 | STM14_5035 | metH | Genic | GA*TC | 3.09 |
| *The *metJ* gene is excluded from these analyses due to artificial excision from the genome |  |  |  |  |  |  |  |  |

### YhdJ plays little role in *Salmonella* physiology under standard conditions important for virulence

While our data do not support a broad, global correlation between differential methylation and differential expression under most of our tested conditions, particularly comparing wild-type bacteria grown in LB and SPI-2 inducing conditions, this does not rule out that there are discreet examples where methylation and gene expression are causally linked in our datasets. We hypothesized that such instances could be identified by combining our data on methylation and transcriptional patterns under biologically relevant conditions with data from methylase knockout mutants to reduce our search space to putative sites of regulation. We tested this hypothesis with the YhdJ (the most dynamic methylase in our dataset) and Dam (the most well-studied DNA methylase in *Salmonella*) mutants.

RNA-seq on wild-type and Δ*yhdJ* bacteria grown in LB or SPI-2 inducing conditions **(GEO: GSE185073, Supplemental File 5)** revealed that knocking out *yhdJ* had almost no impact on the transcriptome. Apart from *yhdJ* itself, only 12 genes were differentially expressed in LB (**Figure 5A, Table 2)** despite loss of methylation at all 513 ATGCA*T sites (see **Supplemental Figure 2C**), and no genes were differentially expressed under SPI-2 inducing conditions **(Figure 5B)**. Curiously, GO-analysis (71,72) demonstrated the differentially expressed genes are enriched for de novo UMP biosynthetic processes (False Discovery Rate=6.3×10^−4^) and de novo pyrimidine nucleobase biosynthetic processes (False Discovery Rate=8.31×10^−4^). However, examining these genes further revealed that only two differentially expressed genes contained or were near an ATGCAT sequence (*dppA* and *pyrB*), and only *dppA* was detected to house a methylated ATGCA*T motif in our Replication Methylation Dataset **(Table 2)**, making the mechanism of this differential expression unclear. In agreement with these findings, *yhdJ* deletion had little impact on cellular or virulence phenotypes. Δ*yhdJ* had no effect on growth in LB or SPI-2 inducing media **(Figure 5C,D)**, a subtle increase on the amount of observed THP-1 cell infection **(Figure 5E)**, no effect on replication in THP-1 cells **(Figure 5F)**, no effect on motility **(Figure 5G)**, and had almost no effect on fitness in intraperitoneal or enteric fever models of mouse infection—though a small increase in the number of Δ*yhdJ* CFUs recovered from the spleen relative to wild-type following oral gavage was observed **(Table 3)**. Across all phenotypes, no genetic interaction between *yhdJ* and *metJ* was detected. Thus, YhdJ has very little impact on transcription or virulence associated phenotypes, and if anything, modestly impairs *Salmonella* virulence.

**Figure 5:**
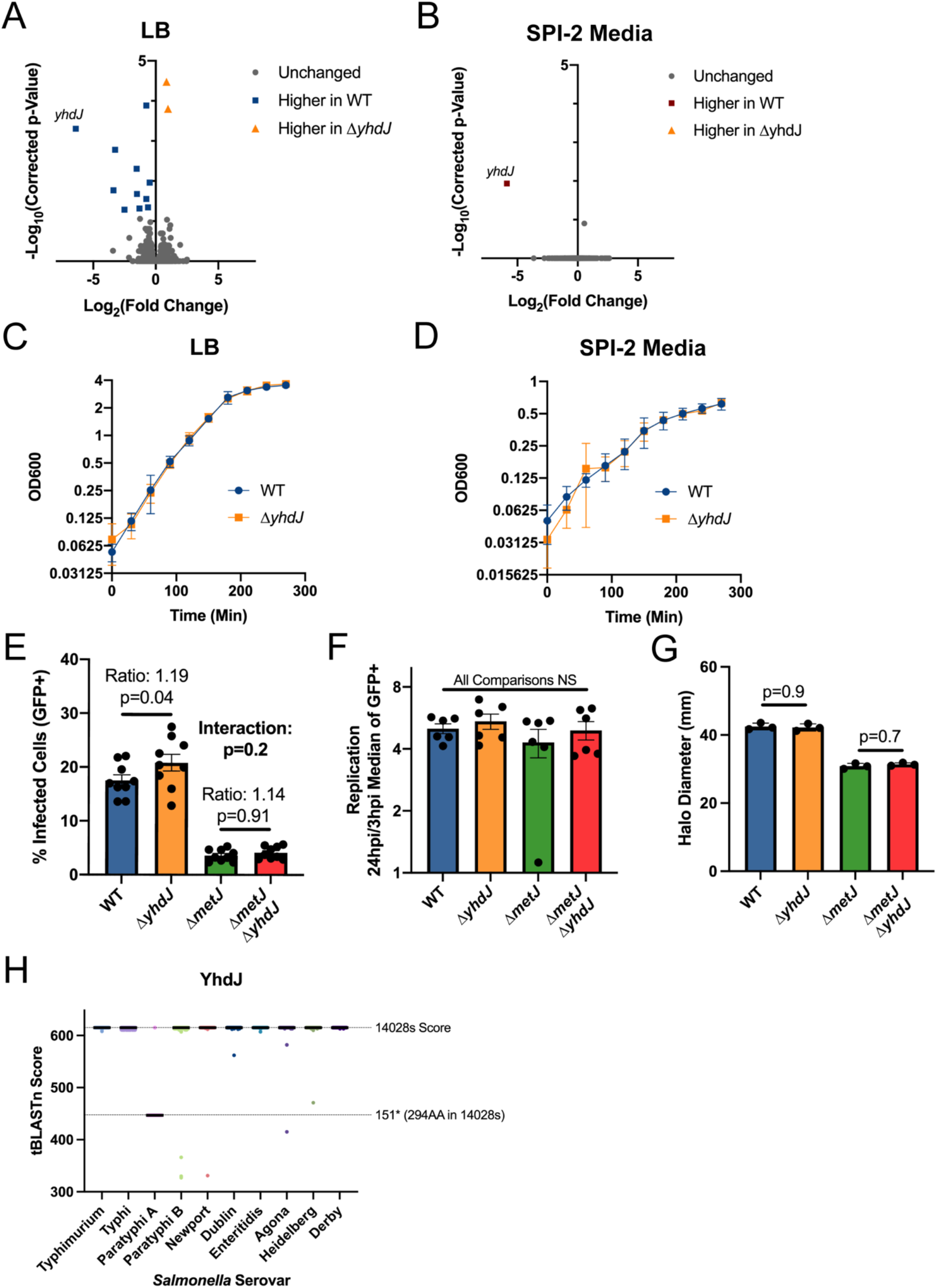
YhdJ has limited impacts on *S*. Typhimurium biology under standard laboratory conditions. (A, B). YhdJ has limited impacts on the *S*. Typhimurium transcriptome in LB (A) and SPI-2 inducing media (B). Corrected p-values generated by calculating the false discovery rate. (C, D) YhdJ is not required for *S*. Typhimurium growth in LB (C) or SPI-2 inducing media (D). Data from three independent experiments with time points taken every 30 minutes. Error bars represent the standard error of the mean. (E, F) YhdJ is not required for *S*. Typhimurium uptake (E) or replication (F) in THP-1 monocytes. Cells were infected for 60 minutes with *S*. Typhimurium harboring an inducible-GFP plasmid before treatment with gentamicin. GFP was induced for 75 minutes before analysis by flow cytometry. Percent GFP+ and median of the GFP+ cells were measured three hours and twenty-four hours post infection. Panel E shows the amount of invasion that occurred by reporting the percent of infected cells at 3 hours post infection, and Panel F shows the replication that occurred following infection by dividing the median of the GFP+ cells at 24 hours post infection by the median of the GFP+ cells at 3 hours post infection. Data from 2-3 independent experiments, each dot represents an independent replicate, bars mark the mean, and error bars are the standard error of the mean. For Panel E, data were normalized to the grand mean before plotting or performing statistics, and for Panel F statistics were performed on the log transformed values. P-values generated by two-way ANOVA with Sidak’s multiple comparisons test. (G) YhdJ does not impact *S*. Typhimurium motility. Motility on soft agar was measured six hours after inoculating the agar and following migration at 37°C. Data are from three independent experiments and each dot is the average of 4-5 technical replicates, bars mark the mean, and error bars mark the standard error of the mean. Data were normalized to the grand mean prior to plotting or performing statistics. P-values generated by one-way ANOVA with Sidak’s multiple comparisons test. (H) YhdJ is conserved across several *Salmonella* serovars. *Salmonella* genomes (1,000 Typhimurium, 1,000 Typhi, 1,000 Paratyphi A, 1,000 Paratyphi B, 999 Newport, 1,000 Dublin, 1,000 Enteritidis, 1,000 Agona, 1,000 Heidelberg, and 79 Derby genomes) were obtained from EnteroBase (68,69). Genomes were combined into a single FASTA file per serovar and blasted against the *S*. Typhimurium strain 14028s YhdJ protein sequence using BLAST+ (70). The BLAST score from the top ‘n’ hits were plotted, where ‘n’ is the number of genomes analyzed for that serovar. Black bar represents the median. Dotted lines represent the BLAST score obtained when blasting the 14028s genome, and the score obtained from the 151* truncation prevalent in *S*. Paratyphi A serovars.

**Table 2:** Δ*yhdJ* Differential Gene Expression in LB.

| Table 2: *ΔyhdJ* Differential Gene Expression in LB |  |  |  |  |  |
| --- | --- | --- | --- | --- | --- |
| Gene ID | Gene Name | Log<sub>2</sub>Fold Change (*ΔyhdJ*/WT) | Corrected p-value | ATGCAT Motif? (Relative Location) | Methylated? |
| STM14_4375 | *dppA* | -0.75 | 2.95E-08 | Yes (Genic) | Yes |
| STM14_4084 | *yhdJ* | -6.39 | 2.25E-07 | No | No |
| STM14_5353 | *pyrB* | -3.25 | 1.12E-06 | Yes (Genic) | No |
| STM14_4024 | *codB* | -1.52 | 4.51E-06 | No | No |
| STM14_0699 | *cstA* | -0.48 | 1.25E-05 | No | No |
| STM14_5352 | *pyrI* | -3.38 | 2.32E-05 | No | No |
| STM14_0819 | *NA* | 0.85 | 3.34E-05 | No | No |
| STM14_4495 | *pyrE* | -1.49 | 3.84E-05 | No | No |
| STM14_5141 | *acs* | -0.74 | 5.74E-05 | No | No |
| STM14_1558 | *yeaG* | -0.62 | 0.0001 | No | No |
| STM14_3061 | *uraA* | -1.3 | 0.0001 | No | No |
| STM14_0078 | *carB* | -2.49 | 0.0001 | No | No |
| STM14_1885 | *hdeB* | 0.99 | 0.0002 | No | No |

**Table 3:** YhdJ does not enhance *S*. Typhimurium fitness in C57BL/6J mice.

| Table 3: YhdJ does not enhance *S. Typhimurium* fitness in C57BL/6J mice |  |  |  |  |
| --- | --- | --- | --- | --- |
| IP Infection (CFUs from Spleen) | $\Delta metJ/WT$ | $\Delta yhdJ/WT$ | $\Delta metJ\Delta yhdJ/WT$ | $\Delta metJ\Delta yhdJ/\Delta metJ$ |
| *Number of Mice*<sup>§</sup> | 6 | 8 | 7 | 7 |
| *Median*<sup>#</sup> (95% Confidence Interval) | **0.23** (0.07-0.47) | **1.31** (0.60-1.84) | **0.26** (0.13-0.78) | **1.09** (0.000065-2.19) |
| *p-value* * | **0.002** | 0.3 | **0.002** | 0.2 |
| Oral Infection (CFUs from Spleen and Ileum) | $\Delta yhdJ/WT$ (Spleen) | $\Delta yhdJ/WT$ (Ileum) | | |
| *Number of Mice*<sup>§</sup> | 22 | 21 |  |  |
| *Median*<sup>#</sup> (95% Confidence Interval) | **1.65** (0.72-2.22) | **1.154** (0.74-1.55) |  |  |
| *p-value* * | **0.02** | 0.4 |  |  |
| <sup>§</sup>All mice are age and sex matched, both sexes represented in all experiments, and all data are from at least two experiments<br><sup>#</sup>Median competitive index value calculated by dividing the number colonies obtained of each genotype at four days post infection. Median > 1 = numerator strain outcompeted denominator.<br>*One-sample t-test on log transformed data |  |  |  |  |

These findings suggest that YhdJ is almost entirely dispensable under the conditions tested here, despite methylating over 500 sites in the genome and methylation of its ATGCA*T motif being the most dynamic under the conditions tested. We questioned whether YhdJ may play roles outside of transcription, and whether its dynamic nature is incidental. Supporting this hypothesis, an evolutionary analysis of over 9,000 strains from 10 serovars revealed that the methylase is highly conserved, with the exception of *S*. Paratyphi A where most strains harbor a 151* truncation **(Figure 5H)**. This conservation paired with our RNA-seq data suggests that YhdJ methylation is likely important but is unlikely to play a large role in gene regulation.

### Integration of previous Δ*dam* literature with methylomics data reveals *stdA* hypomethylation correlates with expression in LB

After demonstrating that a Δ*yhdJ* mutant could not be leveraged to identify instances where gene expression and methylation are linked, we turned to the Δ*dam* literature. Previous work on *Salmonella* transcriptomics revealed 17 virulence genes (making up 11 operons) that are differentially expressed between wild-type and Δ*dam S*. Typhimurium grown in LB to mid-exponential phase (31). We hypothesized that these sites may show differential methylation and expression between our LB and SPI-2 medias, as *Salmonella* deploy radically different virulence programs across these conditions and many of these 17 virulence genes are known to be expressed specifically under only one of these two conditions (14 of 17 were DEGs comparing these two conditions in our data, and the remaining three (STM14_3654, *stdB*, and *stdC*) lacked high enough expression to analyze). In order to test this hypothesis, we searched for GATC motifs upstream (within 500bp) of the DEGs. Interestingly, 10/11 genes or operons we examined contain GATC sites within 500bp upstream of these genes **(Table 4)**. However, only one gene (*stdA*) showed evidence of differential methylation under physiological conditions. Interestingly, *stdA* is the only one of these 11 genes/operons where methylation of its promoter has been extensively studied (58,97). The three GATC sites had reduced methylation following growth in LB, however, each was only hypomethylated on a single strand (**Figure 6A)**. Interestingly, this hypomethylation agrees with a previous report that Dam and transcription factors compete for binding to the *stdA* promoter (58), as we also observed significantly higher expression of *stdA* in LB than in SPI-2 media **(Figure 6B)**. Whether this represents a mechanism by which *stdA* is turned off in SPI-2 media or is a correlated consequence of increased *stdA* transcription in LB (or vice versa) is unclear, but this finding demonstrates that the phenomenon that García-Pastor *et al*. describe in Δ*dam* mutants occurs naturally under biologically relevant conditions. For the other 10 virulence genes/operons that are reported to undergo differential gene expression in the *dam* mutant, we find no evidence that differential methylation plays any role during their induction during SPI-1 or SPI-2-inducing conditions.

**Figure 6:**
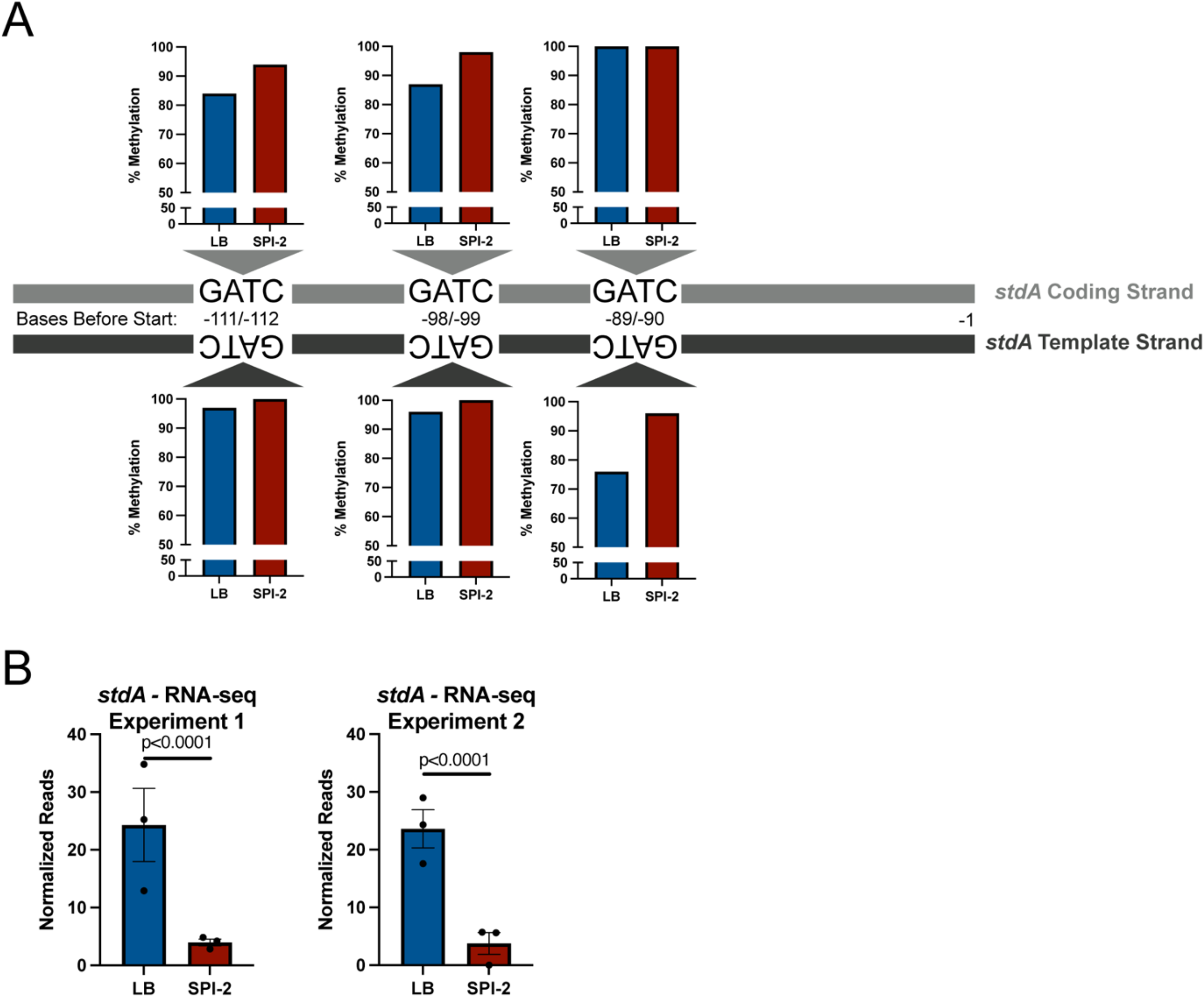
The *stdA* promoter has differential methylation following growth in LB or SPI-2 inducing media. (A) Schematic of the region upstream of *stdA*. Percent methylation values for each GATC site on both strands are graphed based on data from wild-type bacteria in Methylation Experiment 1. (B) *stdA* is differentially expressed in wild-type bacteria grown in LB and SPI-2 inducing media. RNA-seq Experiment 1 values are from the RNA-seq experiment including Δ*metJ* and are listed in **Supplemental File 4**. RNA-seq Experiment 2 values are from the experiment including Δ*yhdJ* and are listed in **Supplemental File 5**.

**Table 4:**
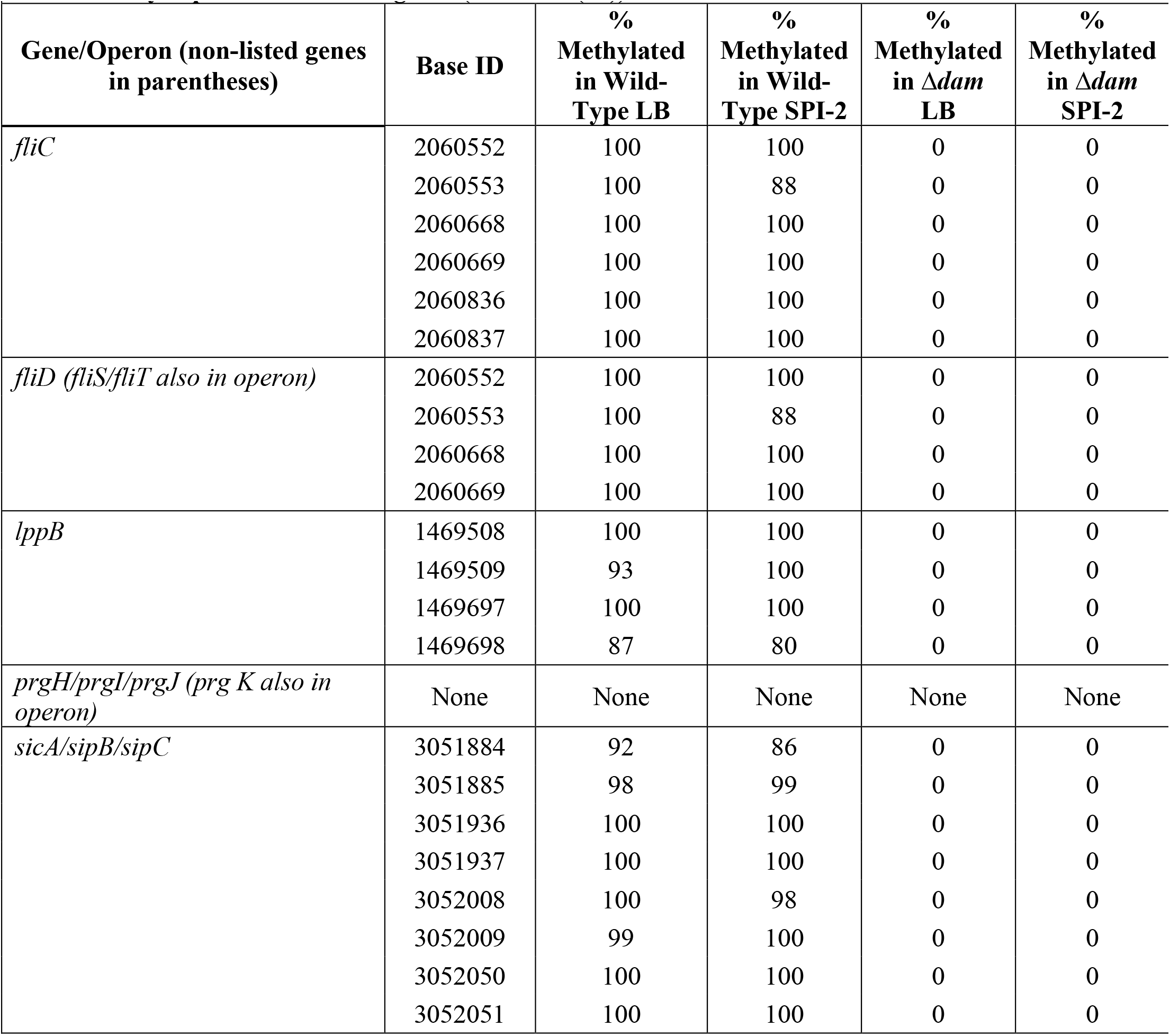
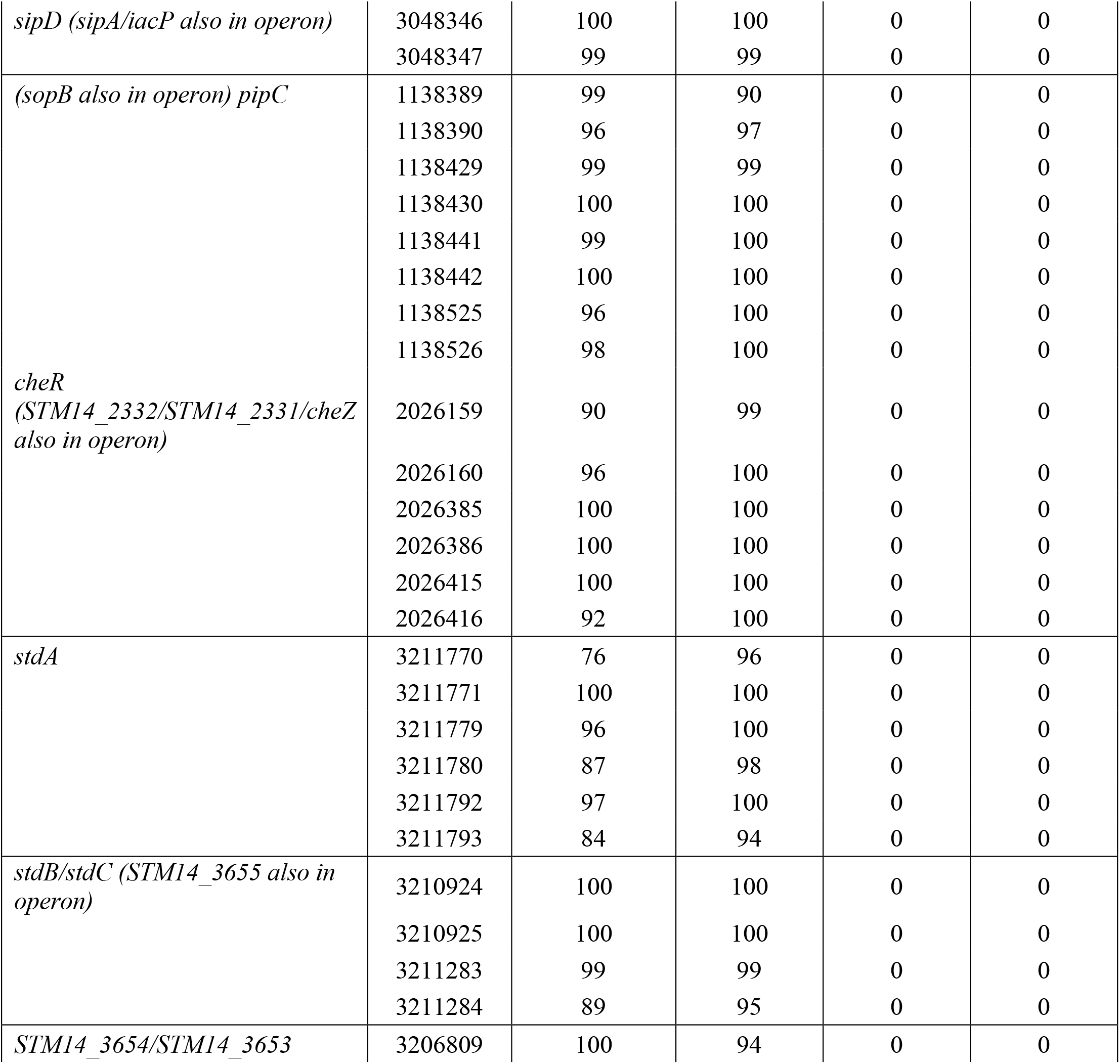
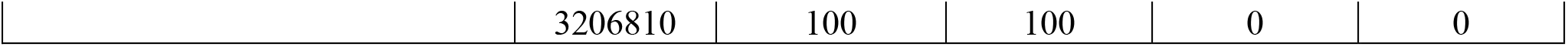
Percent methylation of GATC sites in Experiment 1 within 500bp upstream of Δ*dam* differentially expressed virulence genes (based on (31))

| **Table 4: Percent methylation of GATC sites in Experiment 1 within 500bp upstream of $\Delta dam$ differentially expressed virulence genes (based on (31))** |  |  |  |  |  |
| --- | --- | --- | --- | --- | --- |
| **Gene/Operon (non-listed genes in parentheses)** | **Base ID** | **% Methylated in Wild-Type LB** | **% Methylated in Wild-Type SPI-2** | **% Methylated in $\Delta dam$ LB** | **% Methylated in $\Delta dam$ SPI-2** |
| *fliC* | 2060552 | 100 | 100 | 0 | 0 |
|  | 2060553 | 100 | 88 | 0 | 0 |
|  | 2060668 | 100 | 100 | 0 | 0 |
|  | 2060669 | 100 | 100 | 0 | 0 |
|  | 2060836 | 100 | 100 | 0 | 0 |
|  | 2060837 | 100 | 100 | 0 | 0 |
| *fliD* (*fliS/fliT* also in operon) | 2060552 | 100 | 100 | 0 | 0 |
|  | 2060553 | 100 | 88 | 0 | 0 |
|  | 2060668 | 100 | 100 | 0 | 0 |
|  | 2060669 | 100 | 100 | 0 | 0 |
| *lppB* | 1469508 | 100 | 100 | 0 | 0 |
|  | 1469509 | 93 | 100 | 0 | 0 |
|  | 1469697 | 100 | 100 | 0 | 0 |
|  | 1469698 | 87 | 80 | 0 | 0 |
| *prgH/prgI/prgJ* (*prg K* also in operon) | None | None | None | None | None |
| *sicA/sipB/sipC* | 3051884 | 92 | 86 | 0 | 0 |
|  | 3051885 | 98 | 99 | 0 | 0 |
|  | 3051936 | 100 | 100 | 0 | 0 |
|  | 3051937 | 100 | 100 | 0 | 0 |
|  | 3052008 | 100 | 98 | 0 | 0 |
|  | 3052009 | 99 | 100 | 0 | 0 |
|  | 3052050 | 100 | 100 | 0 | 0 |
|  | 3052051 | 100 | 100 | 0 | 0 |
| *sipD* (*sipA/iacP* also in operon) | 3048346 | 100 | 100 | 0 | 0 |
|  | 3048347 | 99 | 99 | 0 | 0 |
| *(sopB* also in operon) *pipC* | 1138389 | 99 | 90 | 0 | 0 |
|  | 1138390 | 96 | 97 | 0 | 0 |
|  | 1138429 | 99 | 99 | 0 | 0 |
|  | 1138430 | 100 | 100 | 0 | 0 |
|  | 1138441 | 99 | 100 | 0 | 0 |
|  | 1138442 | 100 | 100 | 0 | 0 |
|  | 1138525 | 96 | 100 | 0 | 0 |
|  | 1138526 | 98 | 100 | 0 | 0 |
| *cheR*<br>(*STM14_2332/STM14_2331/cheZ*<br>also in operon) | 2026159 | 90 | 99 | 0 | 0 |
|  | 2026160 | 96 | 100 | 0 | 0 |
|  | 2026385 | 100 | 100 | 0 | 0 |
|  | 2026386 | 100 | 100 | 0 | 0 |
|  | 2026415 | 100 | 100 | 0 | 0 |
|  | 2026416 | 92 | 100 | 0 | 0 |
| *stdA* | 3211770 | 76 | 96 | 0 | 0 |
|  | 3211771 | 100 | 100 | 0 | 0 |
|  | 3211779 | 96 | 100 | 0 | 0 |
|  | 3211780 | 87 | 98 | 0 | 0 |
|  | 3211792 | 97 | 100 | 0 | 0 |
|  | 3211793 | 84 | 94 | 0 | 0 |
| *stdB/stdC* (*STM14_3655* also in operon) | 3210924 | 100 | 100 | 0 | 0 |
|  | 3210925 | 100 | 100 | 0 | 0 |
|  | 3211283 | 99 | 99 | 0 | 0 |
|  | 3211284 | 89 | 95 | 0 | 0 |
| *STM14_3654/STM14_3653* | 3206809 | 100 | 94 | 0 | 0 |
|  | 3206810 | 100 | 100 | 0 | 0 |

We also attempted to integrate our data with a recent study that examined genetic heterogeneity in *S*. Typhimurium (98). We examined the 16 hypomethylated sites they identified to determine whether our dataset could find the same signatures of hypomethylation. Of the seven sites we were able to find in our dataset, four showed signs of hypomethylation (**Supplemental Table 4**). Of these four, two sites upstream of *carA* and *dgoR* showed differential methylation across our conditions, however, no consistent trends were observed with differential gene expression across our two RNA-seq datasets **(Supplemental Figure 8)**. Thus, while we were able to observe an association of gene expression and DNA methylation for one canonical example (*stdA*) during growth in LB vs. SPI-2 inducing conditions, we did not find this to be a generalizable phenomenon for other genes reported to be differentially regulated in the *dam* mutant or hypomethylated in *S*. Typhimurium.

### Despite limited changes to the GA*TC methylome, *metJ* and *dam* interact to influence *S*. Typhimurium invasion and motility

Having successfully used Δ*dam* transcriptomics to identify one biologically meaningful co-occurrence of differential expression and methylation, we next attempted to leverage Δ*dam* mutants to test our hypothesis that aberrant methylation in Δ*metJ* bacteria contribute to the impact of Δ*metJ* on invasion and motility (91). Importantly, we observed a small growth defect in Δ*dam* and Δ*dam*Δ*metJ* bacteria, and so all bacteria used for these experiments were grown an extra 30 minutes prior to infection to standardize the growth phase used. Knocking out *dam* only modestly reduced the impact of Δ*metJ* on invasion and is therefore unlikely to be the primary mechanism by which *metJ* deletion impacts invasion **(Figure 7A)**. In contrast, impairment in motility caused by *metJ* deletion was completely abrogated in Δ*dam* genetic background, suggesting that *dam* and *metJ* impact motility through the same pathway **(Figure 7B)**. Importantly, comparing isogenic strains (wild-type vs Δ*metJ*; Δ*dam* vs Δ*damΔmetJ; Δdam* p*dam* vs *ΔdamΔmetJ* p*dam*) revealed that complementation of *dam* on a low copy number plasmid (pWSK129) restored differences between Δ*dam* and Δ*dam*Δ*metJ* bacteria, though we observed that *dam* complementation itself reduces motility further. This is likely due to modest *dam* overexpression, which has previously been reported to be a more potent inhibitor of *S*. Typhimurium 14028s motility than *dam* deletion (34).

**Figure 7:**
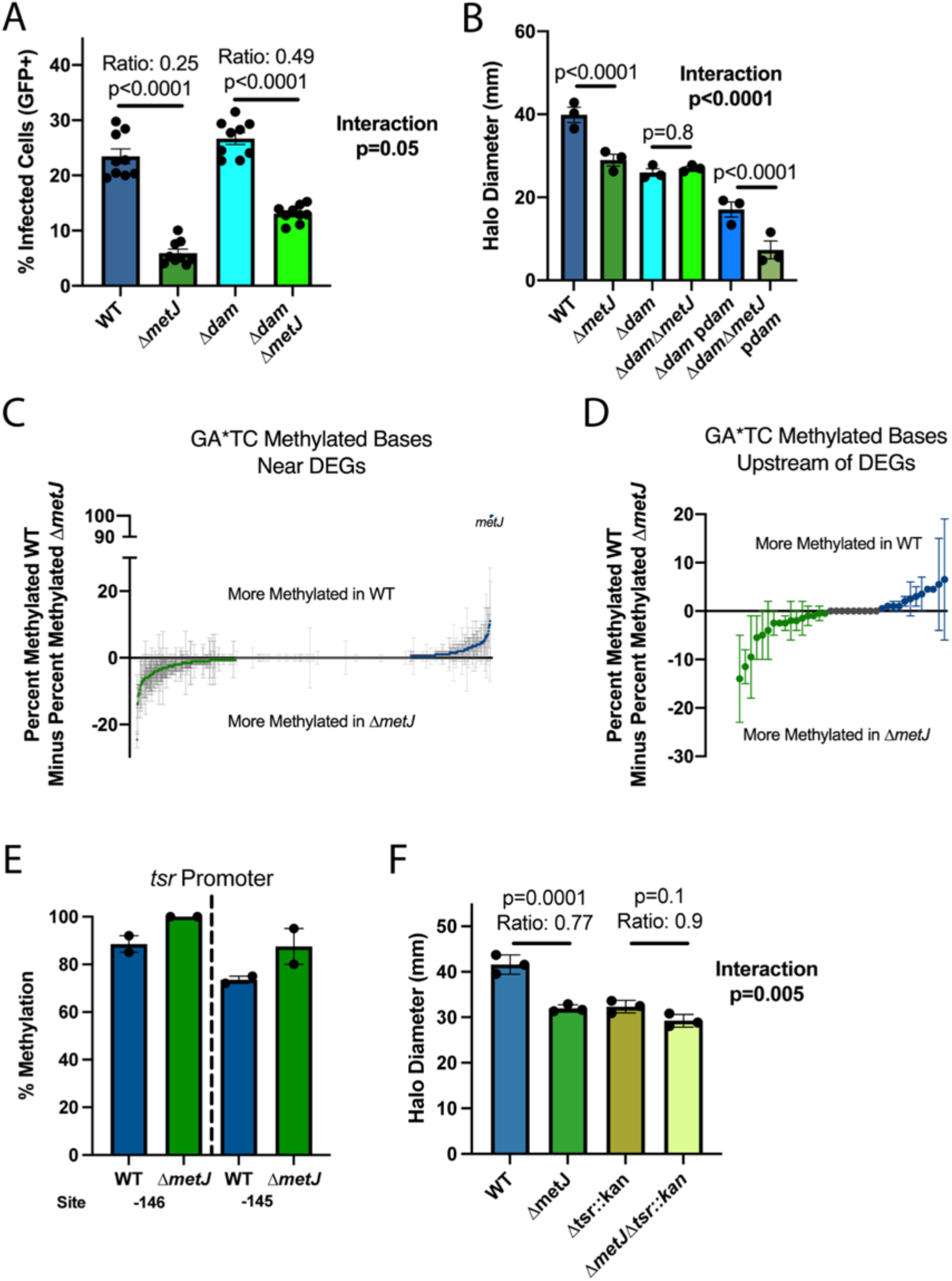
*dam* is epistatic to Δ*metJ* despite limited changes to the Δ*metJ* GA*TC methylome. (A) The impacts of Δ*metJ* on invasion partially depend on *dam*. THP-1 monocytes were infected for 60 minutes with *S*. Typhimurium harboring an inducible-GFP plasmid before treatment with gentamicin. GFP was induced for 75 minutes before analysis by flow cytometry. Percent GFP+ was measured three hours post infection. Data are from three experiments, each dot represents an independent replicate, the bars mark the mean, and the error bars are the standard error of the mean. (B) The impact of Δ*metJ* on motility depends entirely on *dam*. Motility on soft agar was measured six hours after inoculating the agar and following migration at 37°C. Data are from three independent experiments and each dot is the mean of 4-5 technical replicates, bars mark the mean, and error bars mark the standard error of the mean. (C,D) Quantitative analysis reveals subtle changes to the GA*TC methylome in Δ*metJ* bacteria. Each dot represents the difference average percent methylation of GA*TC bases in which the closest gene in differentially expressed (C), or GA*TC bases specifically upstream of differentially expressed genes (D), between WT and Δ*metJ* bacteria grown in LB. Data are in duplicate from the Methylation Experiment 1 and the Replication Methylation Experiment, with error bars showing the error of the mean. Data from Panel D are expanded in **Table 4**. For C and D, any base with greater than or less than 0 differential methylation is colored in green (more methylated in Δ*metJ*) or blue (more methylated in wild-type bacteria). (E) The *tsr* promoter is modestly hypermethylated in Δ*metJ*. Percent methylation is plotted for the -146 and -145 GATC motifs from the Methylation Experiment 1 and the Replication Methylation Experiment, with error bars showing the error of the mean. Site numbering is relative to the start codon. (F) The impacts of Δ*metJ* on motility are partially *tsr* dependent. Data are from three independent experiments and each dot is the mean of 4-5 technical replicates, bars mark the mean, and error bars mark the standard error of the mean. For panels A, B, and F data were normalized to the grand mean prior to plotting or performing statistics and p-values were generated by two-way ANOVA with Sidak’s multiple comparisons test.

### The motility defect of *ΔmetJ S*. Typhimurium partially depends on *tsr*

We hypothesized that the genetic interaction between *dam* and *metJ* could signify that differential GA*TC methylation in the Δ*metJ* mutant suppresses bacterial motility. In striking contrast to this hypothesis, our combined binary dataset revealed no genes that were both differentially GA*TC methylated and expressed (except for the deleted *metJ* itself) **(Table 5)**. We next turned to our percent methylation data to examine whether a shift in methylation could explain differences in flagellar gene regulation between the two bacteria. Comparing percent methylation in both methylation datasets at all GA*TC methylated sites in which the nearest gene is differentially expressed identified 17 sites that had a ≥10% average difference in methylation between wild-type and Δ*metJ* bacteria **(Figure 7C)**. Because co-occurrence of differential methylation and differential expression is expected to occur frequently by chance, we sought to limit our analyses to bases most likely to impact gene expression. To do this, we restricted our search to GA*TC sites that are upstream of differentially expressed genes and found two sites of interest. Specifically, these sites are both strands of a single GATC motif upstream of the chemotaxis gene *tsr* that shows elevated methylation in Δ*metJ* **(Figure 7D and 7E, Table 6)**.

**Table 5:** Unique GA*TC sites between wild-type and Δ*metJ S*. Typhimurium (Combined Dataset)

| Table 5: Unique GA*TC sites between wild-type and $\Delta metJ$ *S. Typhimurium* (Combined Dataset) | | | | | |
| --- | --- | --- | --- | --- | --- |
| Unique WT LB GA*TC Sites |  |  |  |  |  |
| *Genomic Location* | *Base* | *Genic Location* | *Gene* | *Product* | *Differentially Expressed?* |
| None | None | None | None | None | N/A |
| Unique $\Delta metJ$ LB GA*TC Sites | | | | | |
| *Genomic Location* | *Base* | *Genic Location* | *Gene* | *Product* | *Differentially Expressed?* |
| NC_016856.1 | 1177765 | Upstream | STM14_1293 | N-acetylneuraminate Epimerase | No |
| NC_016856.1 | 4289018 | Downstream | STM14_4888 | CDP-diacylglycerol Pyrophosphatase | No |
| *The *metJ* gene is excluded from these analyses due to artificial excision from the genome |  |  |  |  |  |

**Table 6:** Percent methylation differences for GA*TC motifs upstream of Δ*metJ* differentially expressed genes following growth in LB.

| <u>Base</u> | <u>Closest Gene (STM14 Number)</u> | <u>Closest Gene (Common Name)</u> | <u>% Methylation (WT-*ΔmetJ*) Methylation Experiment 1</u> | <u>% Methylation (WT-*ΔmetJ*) Replication Experiment</u> | <u>% Methylation (WT-*ΔmetJ*) Average</u> | <u>Gene Expression - Log<sub>2</sub>Fold Change (*ΔmetJ*/WT)</u> | <u>Gene Expression - Corrected p-value</u> |
| --- | --- | --- | --- | --- | --- | --- | --- |
| 4802770 | STM14 5446 | *tsr* | -23 | -5 | **-14** | -1.28 | 0.006 |
| 4802769 | STM14 5446 | *tsr* | -15 | -8 | **-11.5** | -1.28 | 0.006 |
| 2033749 | STM14 2341 | *flhD* | -1 | -18 | **-9.5** | -0.53 | 0.05 |
| 3243659 | STM14 3699 | *serA* | -10 | -1 | **-5.5** | 1.48 | 5.09E-13 |
| 4415526 | STM14 5029 | *aceB* | -10 | 0 | **-5** | 1.88 | 6.67E-06 |
| 3335887 | STM14 3821 |  | 2 | -10 | **-4** | -1.21 | 0.04 |
| 3400695 | STM14 3893 |  | 0 | -5 | **-2.5** | -1.32 | 0.007 |
| 3400696 | STM14 3893 |  | -2 | -3 | **-2.5** | -1.32 | 0.007 |
| 4415527 | STM14 5029 | *aceB* | -1 | -4 | **-2.5** | 1.88 | 6.67E-06 |
| 2060553 | STM14 2378 |  | 2 | -6 | **-2** | -1.52 | 0.0005 |
| 3335886 | STM14 3821 |  | -4 | 0 | **-2** | -1.21 | 0.04 |
| 571705 | STM14 0600 |  | 2 | -5 | **-1.5** | 4.1 | 1.63E-39 |
| 4415556 | STM14 5029 | *aceB* | -3 | 1 | **-1** | 1.88 | 6.67E-06 |
| 4802790 | STM14 5446 | *tsr* | -1 | -1 | **-1** | -1.28 | 0.006 |
| 3758504 | STM14 4305 |  | -2 | 1 | **-0.5** | -1.41 | 6.13E-06 |
| 3846384 | STM14 4392 |  | -1 | 0 | **-0.5** | 3.05 | 3.03E-07 |
| 571704 | STM14 0600 |  | 0 | 0 | **0** | 4.1 | 1.63E-39 |
| 2060669 | STM14 2378 |  | 0 | 0 | **0** | -1.52 | 0.0005 |
| 2746734 | STM14 3135 | *hmpA* | 0 | 0 | **0** | 1.79 | 7.28E-05 |
| 2746735 | STM14 3135 | *hmpA* | 0 | 0 | **0** | 1.79 | 7.27E-05 |
| 3243658 | STM14 3699 | *serA* | 0 | 0 | **0** | 1.48 | 5.09E-13 |
| 3402626 | STM14 3894 |  | 0 | 0 | **0** | -1.23 | 7.87E-08 |
| 4185969 | STM14 4772 |  | 0 | 0 | **0** | 3.17 | 4.96E-10 |
| 4185987 | STM14 4772 |  | 0 | 0 | **0** | 3.17 | 4.96E-10 |
| 4185988 | STM14 4772 |  | 0 | 0 | **0** | 3.17 | 4.96E-10 |
| 4415557 | STM14 5029 | *aceB* | 1 | 0 | **0.5** | 1.88 | 6.67E-06 |
| 1049501 | STM14 1130 | *ompF* | 1 | 1 | **1** | 1.08 | 4.46E-07 |
| 2060668 | STM14 2378 |  | 2 | 0 | **1** | -1.52 | 0.0005 |
| 4185970 | STM14 4772 |  | 2 | 0 | **1** | 3.17 | 4.96E-10 |
| 3758505 | STM14 4305 |  | 1 | 3 | **2** | -1.41 | 6.13E-06 |
| 3846383 | STM14 4392 |  | -1 | 6 | **2.5** | 3.05 | 3.04E-07 |
| 2060552 | STM14 2378 |  | 5 | 1 | **3** | -1.52 | 0.0005 |
| 4802791 | STM14 5446 | *tsr* | 7 | 0 | **3.5** | -1.28 | 0.006 |
| 1049502 | STM14 1130 | *ompF* | 5 | 4 | **4.5** | 1.08 | 4.46E-07 |
| 3402625 | STM14 3894 |  | 4 | 5 | **4.5** | -1.23 | 7.86E-08 |
| 2072703 | STM14 2394 | *fliJ* | 15 | -4 | **5.5** | -1.55 | 0.006 |
| 2072704 | STM14 2394 | *fliJ* | 19 | -6 | **6.5** | -1.55 | 0.006 |

\*The *metJ* gene is excluded from these analyses due to artificial excision from the genome

This hypermethylation led us to hypothesize that increased methylation upstream of *tsr* in Δ*metJ* could decrease *tsr* expression and thereby reduce motility. In line with this hypothesis, replacing *tsr* with a kanamycin resistance cassette partially ablated the ability for Δ*metJ* to cause a motility defect **(Figure 7F**, interaction term p=0.005). Curiously, a search for the methylation-sensitive transcription factor CRP (41) binding motif (AAATGTGATCTAGATCACATTT) in the *tsr* promoter with the MEME FIMO Tool (99) demonstrated that the hypermethylated residue lies within a putative CRP binding site. Together, these data tentatively support a model in which hypermethylation upstream of *tsr* in Δ*metJ* may contribute to the motility defect. However, additional studies are necessary to confirm a causal relationship.

### Increased Methylation in the *flhDC* promoter does not contribute to the Δ*metJ* motility defect

Given that unlike Δ*dam* (**Figure 7B)**, Δ*tsr* does not account for the entire impact of *metJ* deletion on motility **(Figure 7F**, Ratio of Δ*tsr::kanΔmetJ/*Δ*tsr::kan* = 0.9), we hypothesized that there may be additional differences in GA*TC methylation between wild-type and Δ*metJ* bacteria that impact motility. Further examination of our quantitative methylation dataset **(Table 6)** revealed one additional plausible hypothesis: a site in the *flhDC* promoter (−278) that barely missed our 10% threshold (9.5% more methylated in Δ*metJ* bacteria). We decided to test this site as well, as FlhDC make up the master flagellar regulator and thus modest methylated-mediated regulation of the operon could explain our findings. To test whether differential methylation of the *flhDC* promoter could explain the motility defect in Δ*metJ*, we performed site directed mutagenesis on the *S*. Typhimurium chromosome to mutate the base from GATC to GTTC. However, this mutation had no effect on motility in wild-type or Δ*metJ* bacteria **(Supplemental Figure 9)**, disproving the hypothesis that this site could contribute to the Δ*dam* epistatic effect. Notably, this does not rule out that hypermethylation of this site could play a role in flagellar gene expression in other contexts but does demonstrate that it does not contribute to the Δ*metJ* motility defect.

## Discussion

In this work, we demonstrate that at the genome-wide level differential methylation and differential expression are not correlated. Under the critical conditions of SPI-1 or SPI-2 induction, we observed no association between DEGs and DMGs, whether examining binary changes in methylation or quantitative shifts in methylation of >10%. However, our results do demonstrate that genome-wide methylation studies of biologically relevant conditions can be integrated with data from methylase knockout mutants to identify methyl-bases that may be coupled with gene expression, as exemplified by the *stdA* and *tsr* examples. Integration of data from future methylomic studies with our publicly available datasets could reveal additional naturally occurring instances and potentially important co-occurrence of differential methylation and differential expression. As our work demonstrates, such instances will likely be challenging to identify, as they do not occur more often than expected by chance, and therefore do not appear to be a general mechanism of gene regulation in *Salmonella*. Additionally, we hope that this work encourages the generation of additional methylomic datasets under diverse and biologically relevant conditions in order to enable more intra-species comparative methylomics.

A surprising aspect of our work was that the most differentially active methylase we observed, YhdJ, appears to have almost no impact on the *S*. Typhimurium transcriptome under standard conditions. In contrast to Dam which has known impacts on DNA and bacterial replication (10-19), YhdJ also appears to be completely non-essential for *S*. Typhimurium fitness under our growth conditions and in mice. This raises questions about the broader role of DNA methylation, and in particular YhdJ methylation, in the bacterial cell. One tantalizing hypothesis is that YhdJ plays a role in phage defense, which would have been missed studying the conditions here. Alternatively, YhdJ may contribute to physical genomic structural stability under stress conditions, similar to a proposed role for Dam during antibiotic treatment (32). While these hypotheses could explain why YhdJ does not impact gene expression, they fail to address why we observed reproducible changes in the YhdJ methylome across different conditions. As an answer to this, we speculate that these differences are due to changes in the accessibility of YhdJ to ATGCAT motifs under the different conditions, rather than intentional targeting of YhdJ to these sites. This could be due to differences in other genomic modifications that antagonize YhdJ function, altered protein-DNA interactions that mask ATGCAT sites, and/or changes to the 3D conformation of the genome that prevent interactions between YhdJ and its motif.

We propose three potential explanations for the lack of a consistent correlation between global m^6^A DNA methylation and gene expression in our data. The first is that while *S*. Typhimurium can and do use m^6^A methylation as a mechanism to promote bistability or otherwise regulate transcription, they do so sparingly. This would suggest that while the canonical examples of this are elegant (12,21,22,29,30,36,54-58,98), they are rare exceptions to the general rules of *S*. Typhimurium gene regulation. While we are certainly not the first to discover individual sites of differential m^6^A methylation that do not correlate with gene expression (100,101), this is the first analysis to demonstrate how widespread the phenomenon is in *S*. Typhimurium. The second hypothesis is that three of the four conditions tested here (wild-type or Δ*metJ* bacteria grown in LB or SPI-2 inducing media) are non-representative conditions, whereas our results with the wild-type vs *ΔmetJ* in SPI-2 media are more representative of methylation’s relationship with transcription. Notably, while this is possible, these conditions were specifically chosen as they are (a) relevant to the pathogenic capacity of the bacteria, (b) the conditions most frequently studied in laboratory settings, or (c) disrupt metabolic pathways directly connected to methylation. Therefore, even if methylation plays larger roles in regulating gene expression under other conditions (*e*.*g*. nutrient poor conditions at ambient temperature, following phage insult, etc.), our findings would still suggest that most observed *S*. Typhimurium phenomenon are unlikely to be linked to changes in m^6^A methylation. The third possibility is that while m^6^A is the most common modification to the *S*. Typhimurium genome, other modifications (m^5^C, phosphorothioation, *etc*.) may have more important impacts on gene expression.

In conclusion, through this work we have increased our understanding of the *S*. Typhimurium methylome by defining it as a highly stable system that is largely decoupled from the transcriptome at the genome-wide level. We hope that this work will serve as a reference for how to perform, analyze, and follow-up on DNA methylation studies, and that it will help redefine how we think about m^6^A methylation in bacteria.

## Data Availability

All sequencing data is available in the NCBI’s Gene Expression Omnibus (GEO) (102) Super Series (GSE185077). This includes both SMRT-seq experiments (GSE185578 and GSE185501), as well as both RNA-seq experiments (GSE185072 and GSE185073). All biological resources are available upon request to Dr. Dennis Ko.

## Acknowledgements

The authors would like to thank the Duke University School of Medicine for the use of the Sequencing and Genomic Technologies Shared Resource for performing the library preparations and sequencing experiments referenced throughout this paper. We thank Dr. David Corcoran for supervising the genomic analyses performed by J.L.M and W.C. We thank Kristin Cleveland and Duke DLAR Breeding Core personnel for breeding and maintenance of mouse lines. pREDTKI (Addgene plasmid # 51628 ; http://n2t.net/addgene:51628 ; RRID:Addgene_51628), pMDIAI (Addgene plasmid # 51655 ; http://n2t.net/addgene:51655 ; RRID:Addgene_51655), and pKSI-1(Addgene plasmid # 51725 ; http://n2t.net/addgene:51725 ; RRID:Addgene_51725) were gifts from Sheng Yang. We also thank all past and present members of the Ko lab, especially Kyle Gibbs, Alejandro Antonia, Alyson Barnes, Rachel Keener, Angela Jones, and Margaret Gaggioli for their helpful discussions about the manuscript. Finally, we are grateful to Dr. Stacy Horner and the Duke Molecular Genetics and Microbiology department for use of equipment and shared resources. All schematic images were generated using Biorender.com.

## Funding

This work was supported by the National Institutes of Health [1F31AI143147 to JSB, R01AI118903 to DCK, R21AI144586 to DCK]. The funders played no role in the study design, data collection and analysis, decision to publish, or preparation of the manuscript.

## Conflict of Interest

No authors report a conflict of interest.

## Figures and Tables

**Figure S1: Analysis of changes to the *S*. Typhimurium m**^**6**^**A methylome in response to changing conditions reveals YhdJ as a dynamic methylase**. (A-C) Identification of motifs enriched in methylation sites unique to each of the comparisons in **Figure 2**. Motif enrichment was calculated by dividing the frequency of the motif among the uniquely methylated bases by the genome-wide frequency of that motif within that condition (*ex*. For Panel A, frequency of ATGCA*T within unique WT SPI-2 sites = 242 ATGCA*T sites/423 unique SPI-2 sites (0.57); frequency of ATGCA*T within all WT SPI-2 = 600 ATGCA*T sites/38,843 detected motifs (0.015); enrichment = 0.57/0.015 = 37.04). For all panels, only bases that could be confidently called methylated or unmethylated in all eight conditions in Methylation Experiment 1 were considered.

**Figure S2: A replication screen reveals methylation is highly reproducible across SMRT-seq experiments but highlights the value of performing biological replicates**. (A) Schematic for the Replication Methylation Experiment. Wild-type *S*. Typhimurium (Strain 14028s) or isogenic mutants were grown in LB media and DNA was harvested for SMRT-sequencing. (B) Approximately 97% of bases were called identically (methylated or unmethylated) in Methylation Experiment 1 and the Replicate Methylation Experiments. (C) Only ATGCA*T and “other” sites (bases that do not map to one of the six motifs) change dramatically across tested conditions in the Replication Methylation Experiment. No ATGCA*T methylation was observed in Δ*yhdJ* mutants. (D) The observed Percent Methylation at each base is reproducible across experiments. The color of the hexagon represents the number of bases that fall at that point on the axes. R^2^ values and trendlines represent the correlation across experiments. (E) Quantitative analysis reveals numerous sites are differentially methylated between wild-type and Δ*metJ*. Each dot represents the mean percent methylation in wild-type bacteria across the two experiments subtracted by the mean methylation in Δ*metJ* bacteria for each adenosine confidently called in both experiments. Blue and green dots mark bases where the mean difference is ≥10%. (F) Quantification of unique methylation sites in the Replication Experiment. For Panels C-F, bases were only included in the analysis if the base could confidently be called methylated or unmethylated across conditions. (G) Venn diagram is based on binary measures of differential methylation in the combined dataset. Sites identified by the binary analysis were examined in our quantitative dataset in order to identify changes in the percent methylation. In the graphs “Total” refers to all differentially methylated sites under that condition, and differentially methylated sites are then broken down by motif. For motifs where no differentially methylated sites were present, a single dot is listed at 0%. For shared sites, the absolute value of the difference between bases are shown and thus the numbers are agnostic to whether methylation is higher in either condition. Bars mark the median.

**Figure S3: ACCWGG is enriched in “other” differentially methylated sites**. (A,B) The 40 base pairs flanking sites that were differentially methylated between wild-type and Δ*metJ* bacteria grown in LB in our Combined Dataset (A) or between wild-type bacteria grown in LB and SPI-2 inducing conditions in Methylation Experiment 1 (B) but did not map to one of our 6 motifs were plugged into the MEME software (95) in order to identify overrepresented motifs. ACCWGG, a common miscall for the m^5^C motif CCWGG, was identified in both comparisons.

**Figure S4: Conditions in the RNA-seq experiments cluster with previously published datasets**. PCA analysis comparing data from the Δ*metJ* RNA-seq experiment (Supplemental File 4; “Exp 1”) and the Δ*yhdJ* experiment (Supplemental File 5; “Exp 2”) cluster with data from Kröger *et al*. (96). The LB condition used from Kröger *et al*. was early stationary phase, for which the OD600 (∼2.0) most closely matches the OD600 used in this study (1.5-2.0). Both the SPI-2 inducing condition and the SPI-2+MgCl2 condition were included from Kröger *et al*. Medias separate along PC1, which accounts for 53.5% of the variation. Each condition from this paper is completed in triplicate (with three dots represented on the plot), each Kröger condition in duplicate (two dots on the plot).

**Figure S5: *flhDC* expression is reduced in Δ*metJ***. (A) In contrast to our previous findings (91), *flhD* expression is reduced in Δ*metJ* bacteria by qPCR. Bacteria in late log phase growth in LB were harvested, RNA was stabilized, and RNA was extracted and quantified as described in the methods. Fold change (Δ*metJ*/wild-type) is expressed as 2^-ΔΔCT^, where *flhD* transcript was normalized to the *rrs* gene. Each dot represents the average of 2-3 technical replicates. (B) Endogenous tagging of *flhC* confirms reduced *flhDC* expression in Δ*metJ* bacteria. A C-terminal 3xFLAG tag was added to the *flhC* gene, and abundance of the protein was measured by western blotting. Specificity of the FLAG antibody to FlhC-3xFLAG was confirmed by comparing to a wild-type, untagged *S*. Typhimurium strain. FlhC-3xFLAG abundance was quantified following correcting for loading by normalizing to total protein, and data are presented relative to wild-type (WT) *flhC-3xFLAG S*. Typhimurium. Each dot represents an independent experiment, bars represent the mean, and error bars the standard error of the mean. For A and B, p-values are from a one sample t-test performed on the log transformed data comparing the log(values) to 0.

**Figure S6: Stratification of binary data does not reveal correlation between differentially expressed genes and differentially methylated genes**. (A-C) Fisher’s Exact Test does not reveal an association between differential expression and methylation when the statistical cutoff for differential expression is changed to log2FC>1.5 and FDR corrected p-value<0.05 for any condition. For wild-type LB vs SPI-2 comparisons, there are also no statistical associations when (D) the statistical cutoff for differential expression is changed to log2FC>2.0, or when the data are stratified by (E) direction of gene expression change in LB, (F) direction of gene expression change in SPI-2 media, (G) differentially methylated bases upstream of genes, (H) differentially ATGCA*T methylation, (I) differential GA*TC methylation. Uniquely methylated genes are plotted in the condition under which they are methylated (*e*.*g*. for panel B, a gene that has a methylated upstream base in LB but not SPI-2 media would be plotted as part of “LB”), but are agnostic to the direction of effect for the expression change except in Panels E and F. Data for Panel B from the combined dataset, all other data from Methylation Experiment 1.

**Figure S7: Stratification of quantitative data does not reveal additional correlations between differentially expressed genes and differentially methylated genes**. (A-C) Fisher’s Exact Test does not reveal an association between differential expression and methylation when the statistical cutoff for differential expression is changed to log2FC>1.5 and FDR corrected p-value <0.05 for any condition, except for wild-type bacteria with increased methylation relative to Δ*metJ* bacteria in SPI-2 media where an association was also seen with the less stringent cutoff. (D-F) Fisher’s Exact Test does not reveal an association between differential expression and methylation differentially methylated bases when only differential methylation upstream of genes is considered. (G-I) Fisher’s Exact Test does not reveal an association between differential expression and methylation when the definition of “Differential Methylation” is shifted to sites where (1) the base is ≥99% methylated in one condition, and (2) has a difference of ≥10% between the two conditions, except for wild-type bacteria with increased methylation relative to Δ*metJ* bacteria in SPI-2 media where an association was also seen with the less stringent cutoff. Differentially methylated genes are plotted in the condition under which they are hypermethylated (*e*.*g*. for panel D, a gene that has an upstream base with increased methylation in LB but not SPI-2 media would be plotted as part of “LB”), but are agnostic to the direction of effect for the expression change. Data for panels B, E, and H from the combined dataset, all other data from Methylation Experiment 1.

**Figure S8: Differentially methylated sites from Sánchez-Romero *et al* (98) do not correlate with reproducible changes in gene expression**. (A,B) *carA* (A) and *dgoR* (B) do not show reproducible changes in gene expression between LB grown and SPI-2 induced bacteria. (C,D) Other genes have reproducible effects in the dataset including (C) *prgH* and (D) *ssaT*. Exp 1 refers to the Δ*metJ* RNA-seq experiment (Supplemental File 4) and Exp 2 refers to the Δ*yhdJ* RNA-seq experiment (Supplemental File 5). P-value calculated from FDR.

**Figure S9: Methylation upstream of *flhDC* does not contribute to the Δ*metJ* motility defect**. The -278 GATC sequence is not required for the impacts of Δ*metJ* on motility. Motility on soft agar was measured six hours after inoculating the agar and following migration at 37°C, each dot represents the average of 3-5 technical replicates, data were normalized to the grand mean prior to plotting or performing statistics, and p-values were generated by two-way ANOVA with Sidak’s multiple comparisons test.

**Table S1: Bacterial strains used in this study**

**Table S2: Plasmids used in this study**

**Table S3: Oligonucleotides used in this study**

**Table S4: Percent methylation compared to previous hypomethylation studies**

## Notes

### Competing Interest Statement

The authors have declared no competing interest.

