## Supplemental Figures and Tables for "Integration of the *Salmonella* Typhimurium methylome and transcriptome reveals DNA methylation and transcriptional regulation are largely decoupled under virulence-related conditions"

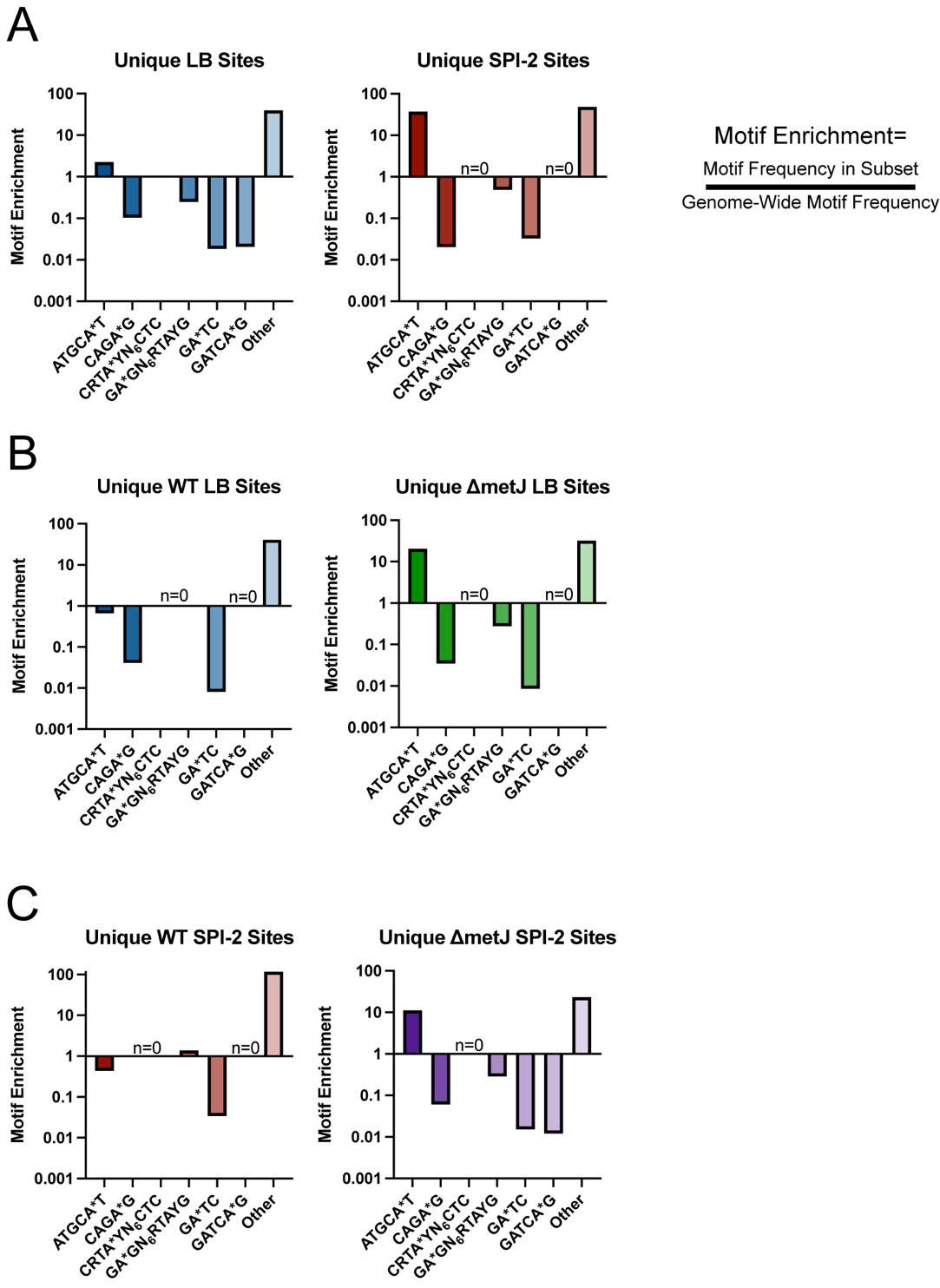

**Supplemental Figure 1: Analysis of changes to the *S.* Typhimurium m^6^A methylome in response to changing conditions reveals YhdJ as a dynamic methylase.** (A-C) Identification of motifs enriched in methylation sites unique to each of the comparisons in **Figure 2**. Motif enrichment was calculated by dividing the frequency of the motif among the uniquely methylated bases by the genome-wide frequency of that motif within that condition (*ex.* For Panel A, frequency of ATGCA*T within unique WT SPI-2 sites = 242 ATGCA*T sites/423 unique SPI-2 sites (0.57); frequency of ATGCA*T within all WT SPI-2 = 600 ATGCA*T sites/38,843 detected motifs (0.015); enrichment = 0.57/0.015 = 37.04). For all panels, only bases that could be confidently called methylated or unmethylated in all eight conditions in Methylation Experiment 1 were considered.

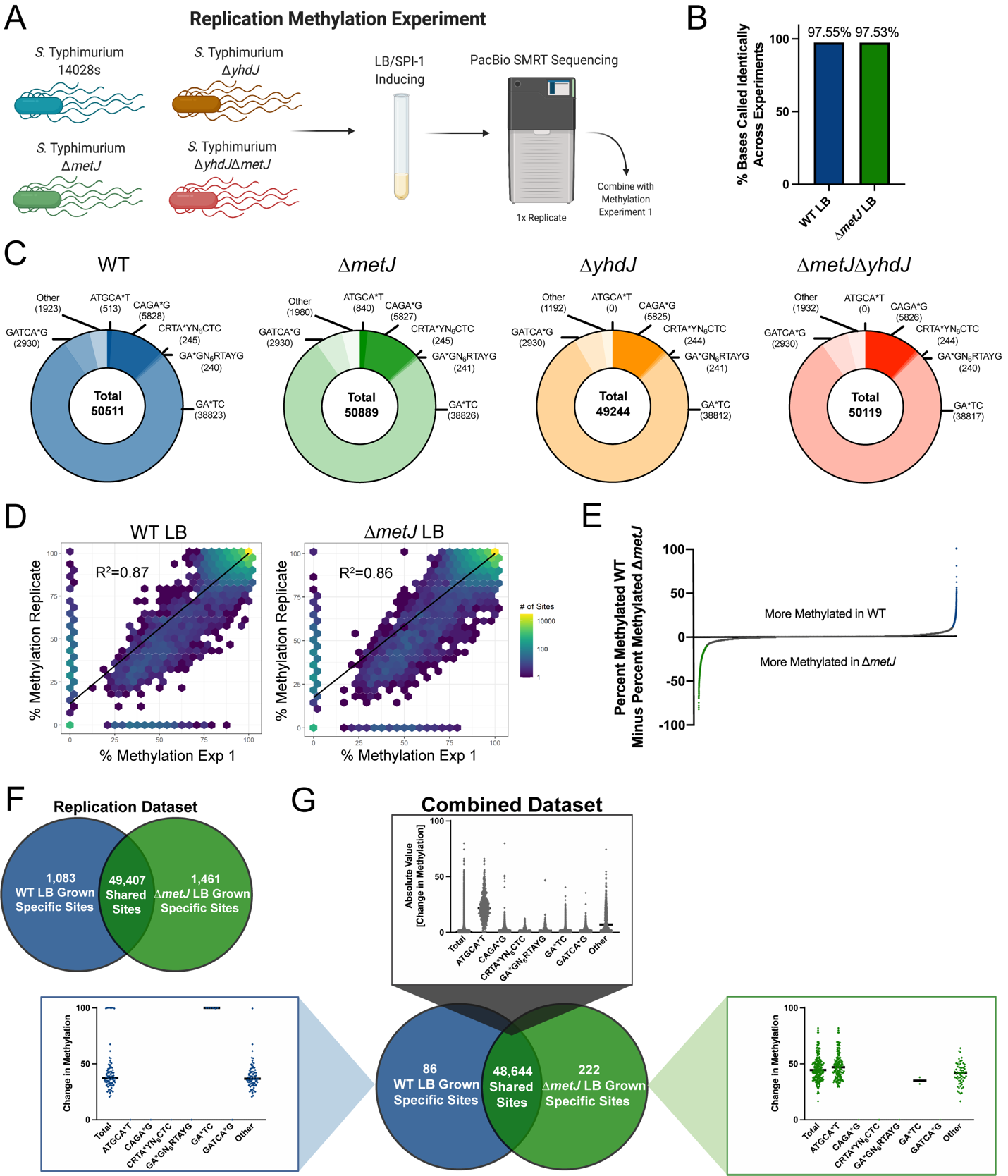

**Supplemental Figure 2: A replication screen reveals methylation is highly reproducible across SMRT-seq experiments but highlights the value of performing biological replicates.** (A) Schematic for the Replication Methylation Experiment. Wild-type *S.* Typhimurium (Strain 14028s) or isogenic mutants were grown in LB media and DNA was harvested for SMRT-sequencing. (B) Approximately 97% of bases were called identically (methylated or unmethylated) in Methylation Experiment 1 and the Replicate Methylation Experiments. (C) Only ATGCA*T and “other” sites (bases that do not map to one of the six motifs) change dramatically across tested conditions in the Replication Methylation Experiment. No ATGCA*T methylation was observed in ∆*yhdJ* mutants. (D) The observed Percent Methylation at each base is reproducible across experiments. The color of the hexagon represents the number of bases that fall at that point on the axes. R^2^ values and trendlines represent the correlation across experiments. (E) Quantitative analysis reveals numerous sites are differentially methylated between wild-type and ∆*metJ*. Each dot represents the mean percent methylation in wild-type bacteria across the two experiments subtracted by the mean methylation in ∆*metJ* bacteria for each adenosine confidently called in both experiments. Blue and green dots mark bases where the mean difference is ≥10%. (F) Quantification of unique methylation sites in the Replication Experiment. For Panels C-F, bases were only included in the analysis if the base could confidently be called methylated or unmethylated across conditions. (G) Venn diagram is based on binary measures of differential methylation in the combined dataset. Sites identified by the binary analysis were examined in our quantitative dataset in order to identify changes in the percent methylation. In the graphs “Total” refers to all differentially methylated sites under that condition, and differentially methylated sites are then broken down by motif. For motifs where no differentially methylated sites were present, a single dot is listed at 0%. For shared sites, the absolute value of the difference between bases are shown and thus the numbers are agnostic to whether methylation is higher in either condition. Bars mark the median.

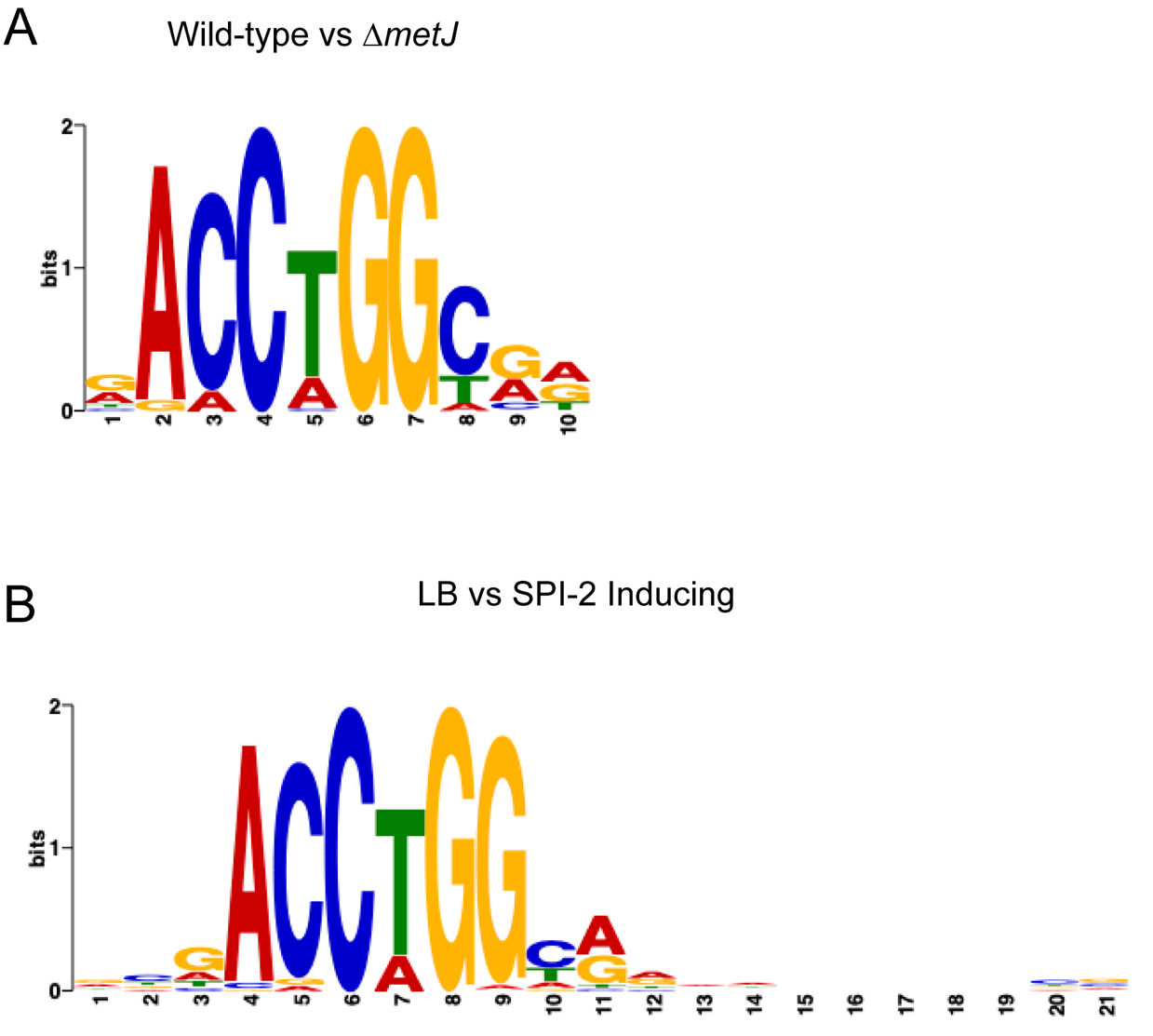

**Supplemental Figure 3: ACCWGG is enriched in “other” differentially methylated sites.** (A,B) The 40 base pairs flanking sites that were differentially methylated between wild-type and ∆*metJ* bacteria grown in LB in our Combined Dataset (A) or between wild-type bacteria grown in LB and SPI-2 inducing conditions in Methylation Experiment 1 (B) but did not map to one of our 6 motifs were plugged into the MEME software (1) in order to identify overrepresented motifs. ACCWGG, a common miscall for the m^5^C motif CCWGG, was identified in both comparisons.

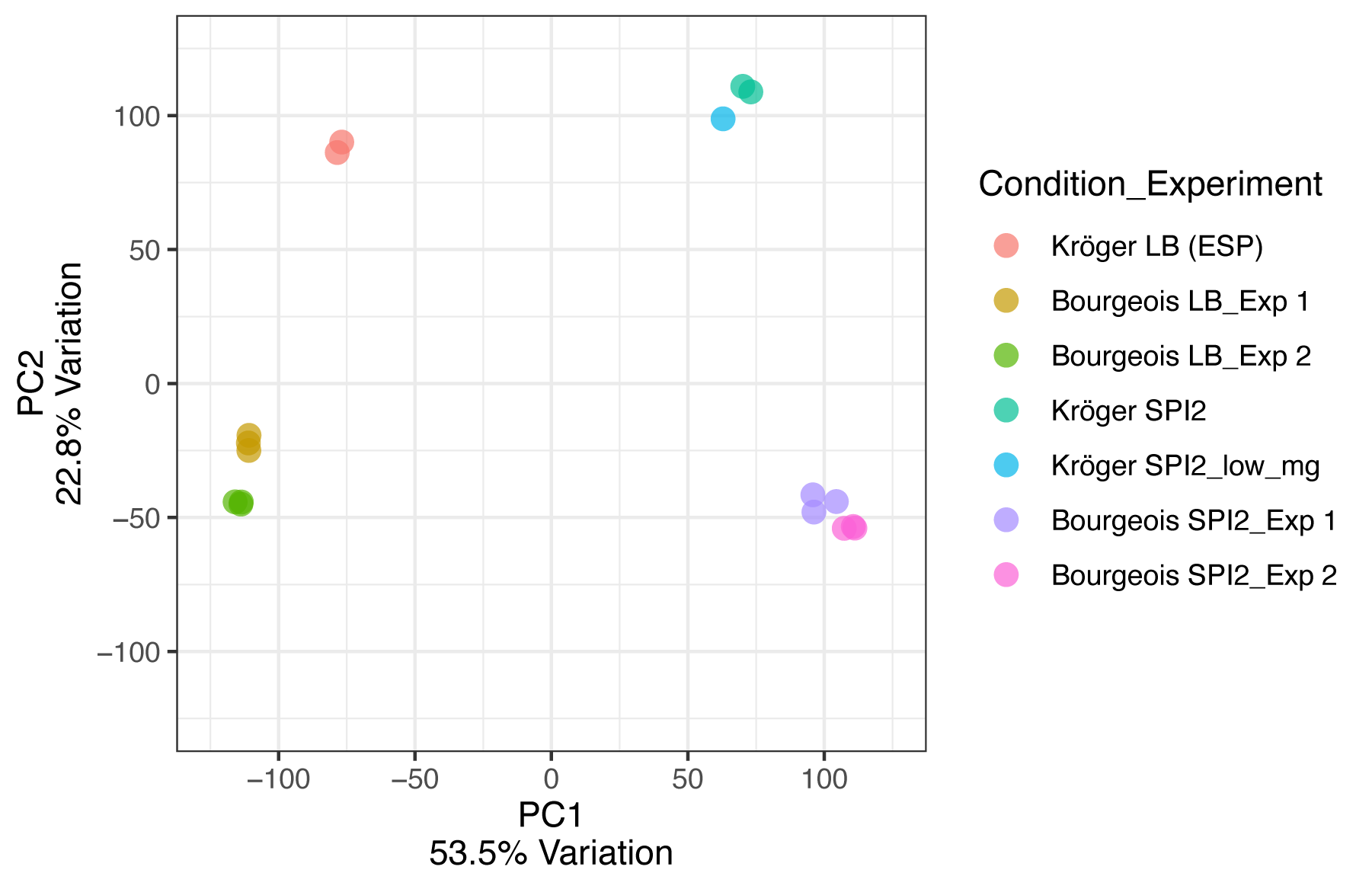

**Supplemental Figure 4: Conditions in the RNA-seq experiments cluster with previously published datasets.** PCA analysis comparing data from the ∆*metJ* RNA-seq experiment (Supplemental File 4; “Exp 1”) and the ∆*yhdJ* experiment (Supplemental File 5; “Exp 2”) cluster with data from Kröger *et al*. (2). The LB condition used from Kröger *et al*. was early stationary phase, for which the OD600 (~2.0) most closely matches the OD600 used in this study (1.5-2.0). Both the SPI-2 inducing condition and the SPI-2+MgCl2 condition were included from Kröger *et al*. Medias separate along PC1, which accounts for 53.5% of the variation. Each condition from this paper is completed in triplicate (with three dots represented on the plot), each Kröger condition in duplicate (two dots on the plot).

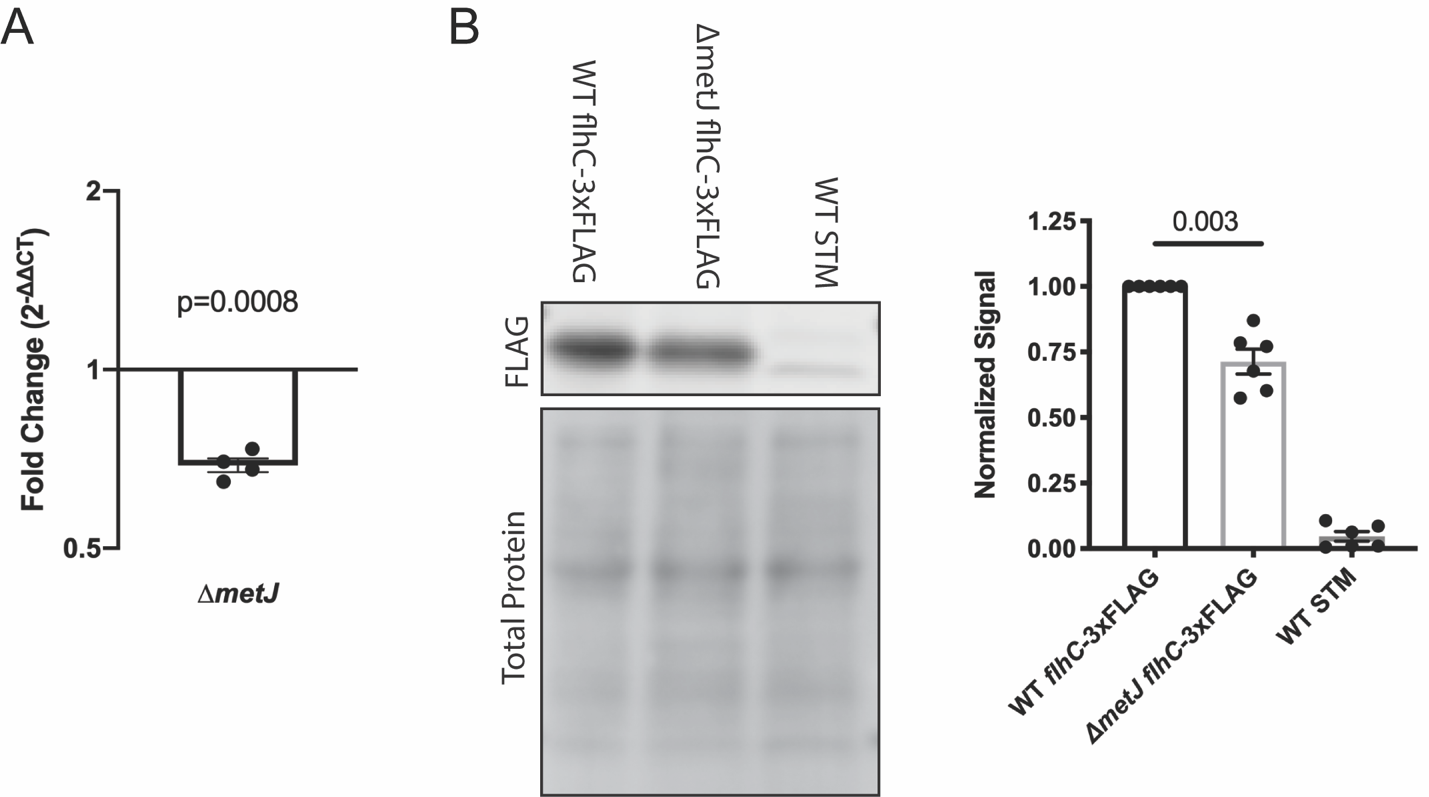

**Supplemental Figure 5: *flhDC* expression is reduced in ∆*metJ*.** (A) In contrast to our previous findings (3), *flhD* expression is reduced in ∆*metJ* bacteria by qPCR. Bacteria in late log phase growth in LB were harvested, RNA was stabilized, and RNA was extracted and quantified as described in the methods. Fold change (∆*metJ*/wild-type) is expressed as 2^-∆∆CT^, where *flhD* transcript was normalized to the *rrs* gene. Each dot represents the average of 2-3 technical replicates. (B) Endogenous tagging of *flhC* confirms reduced *flhDC* expression in ∆*metJ* bacteria. A C-terminal 3xFLAG tag was added to the *flhC* gene, and abundance of the protein was measured by western blotting. Specificity of the FLAG antibody to FlhC-3xFLAG was confirmed by comparing to a wild-type, untagged *S.* Typhimurium strain. FlhC-3xFLAG abundance was quantified following correcting for loading by normalizing to total protein, and data are presented relative to wild-type (WT) *flhC-3xFLAG* *S.* Typhimurium. Each dot represents an independent experiment, bars represent the mean, and error bars the standard error of the mean. For A and B, p-values are from a one sample t-test performed on the log transformed data comparing the log(values) to 0.

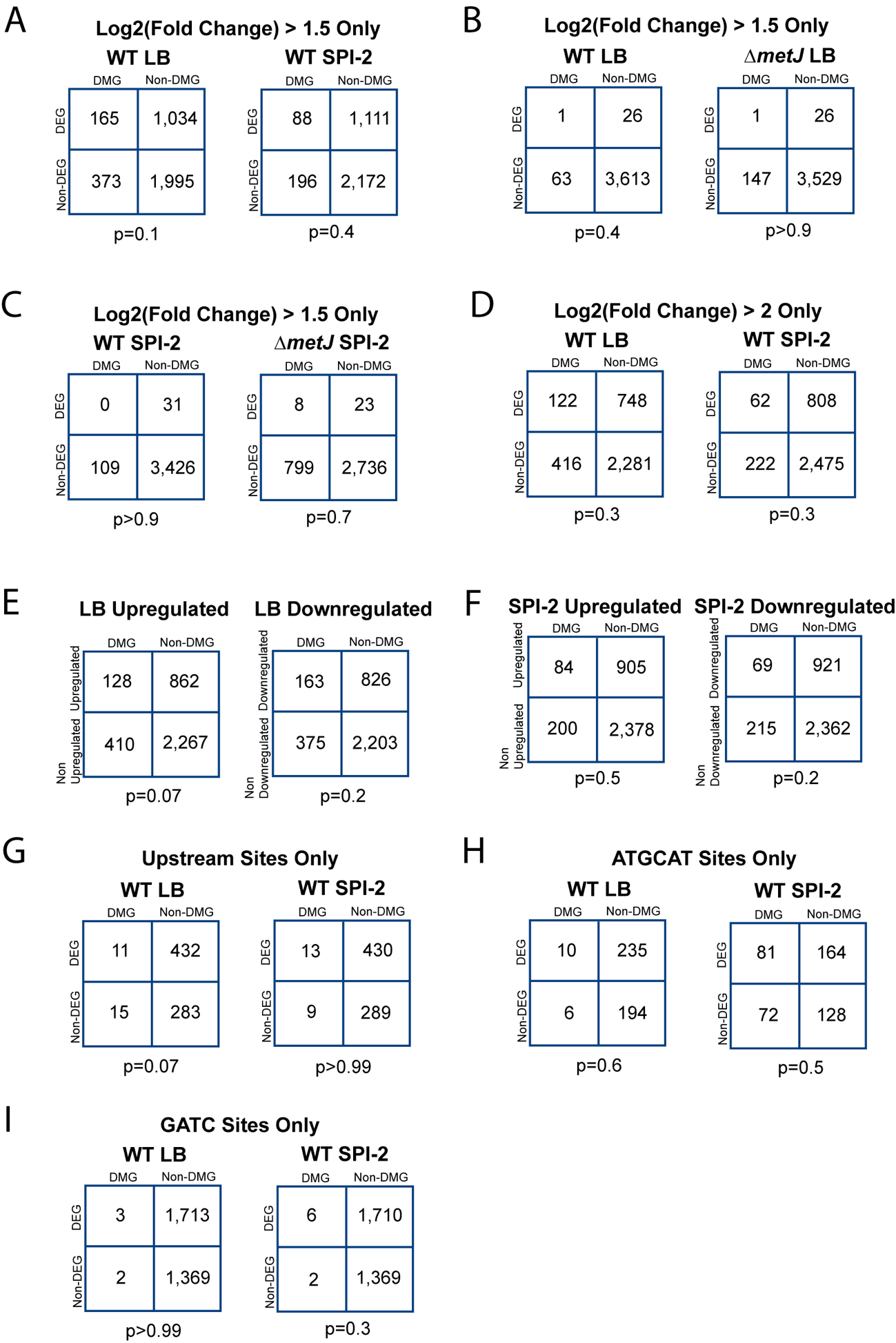

**Supplemental Figure 6: Stratification of binary data does not reveal correlation between differentially expressed genes and differentially methylated genes.** (A-C) Fisher’s Exact Test does not reveal an association between differential expression and methylation when the statistical cutoff for differential expression is changed to log_2_FC>1.5 and FDR corrected p-value<0.05 for any condition. For wild-type LB vs SPI-2 comparisons, there are also no statistical associations when (D) the statistical cutoff for differential expression is changed to log_2_FC>2.0, or when the data are stratified by (E) direction of gene expression change in LB, (F) direction of gene expression change in SPI-2 media, (G) differentially methylated bases upstream of genes, (H) differentially ATGCA*T methylation, (I) differential GA*TC methylation. Uniquely methylated genes are plotted in the condition under which they are methylated (*e.g.* for panel B, a gene that has a methylated upstream base in LB but not SPI-2 media would be plotted as part of “LB”), but are agnostic to the direction of effect for the expression change except in Panels E and F. Data for Panel B from the combined dataset, all other data from Methylation Experiment 1.

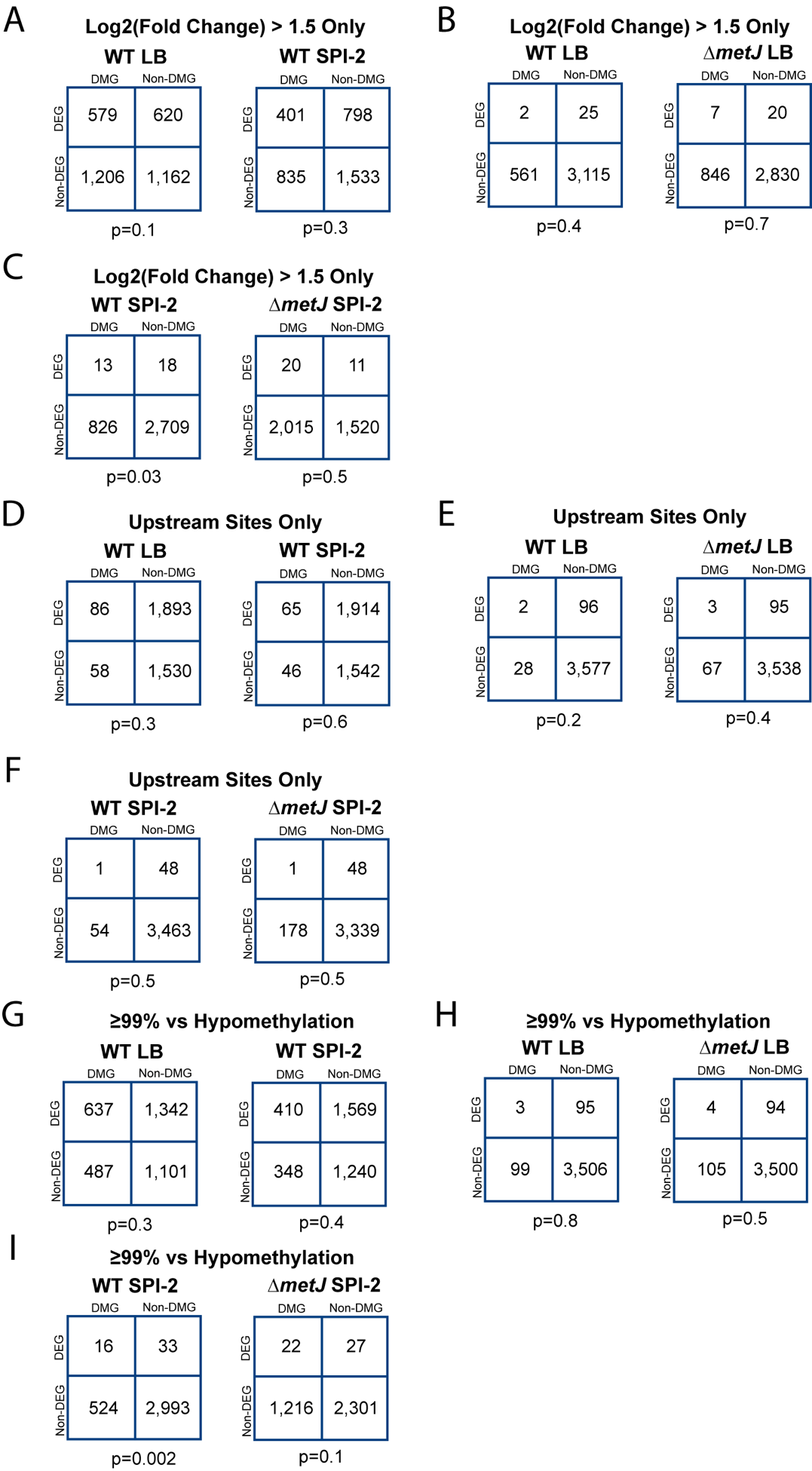

**Supplemental Figure 7: Stratification of quantitative data does not reveal additional correlations between differentially expressed genes and differentially methylated genes.** (A-C) Fisher’s Exact Test does not reveal an association between differential expression and methylation when the statistical cutoff for differential expression is changed to log_2_FC>1.5 and FDR corrected p-value <0.05 for any condition, except for wild-type bacteria with increased methylation relative to ∆*metJ* bacteria in SPI-2 media where an association was also seen with the less stringent cutoff. (D-F) Fisher’s Exact Test does not reveal an association between differential expression and methylation differentially methylated bases when only differential methylation upstream of genes is considered. (G-I) Fisher’s Exact Test does not reveal an association between differential expression and methylation when the definition of “Differential Methylation” is shifted to sites where (1) the base is ≥99% methylated in one condition, and (2) has a difference of ≥10% between the two conditions, except for wild-type bacteria with increased methylation relative to ∆*metJ* bacteria in SPI-2 media where an association was also seen with the less stringent cutoff. Differentially methylated genes are plotted in the condition under which they are hypermethylated (*e.g.* for panel D, a gene that has an upstream base with increased methylation in LB but not SPI-2 media would be plotted as part of “LB”), but are agnostic to the direction of effect for the expression change. Data for panels B, E, and H from the combined dataset, all other data from Methylation Experiment 1.

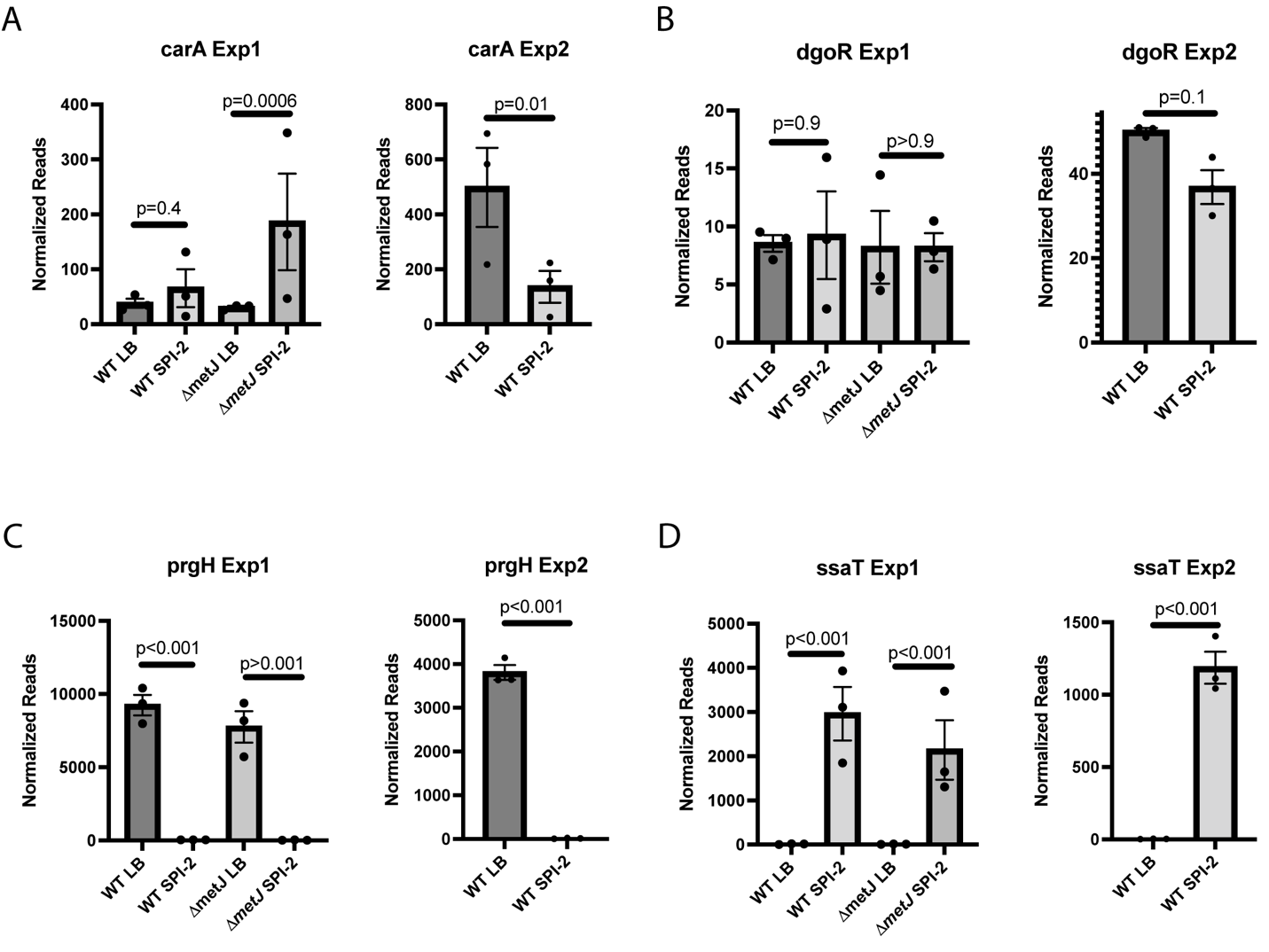

**Supplemental Figure 8: Differentially methylated sites from Sánchez-Romero *et al* (4) do not correlate with reproducible changes in gene expression.** (A,B) *carA* (A) and *dgoR* (B) do not show reproducible changes in gene expression between LB grown and SPI-2 induced bacteria. (C,D) Other genes have reproducible effects in the dataset including (C) *prgH* and (D) *ssaT.* Exp 1 refers to the ∆*metJ* RNA-seq experiment (Supplemental File 4) and Exp 2 refers to the ∆*yhdJ* RNA-seq experiment (Supplemental File 5). P-value calculated from FDR.

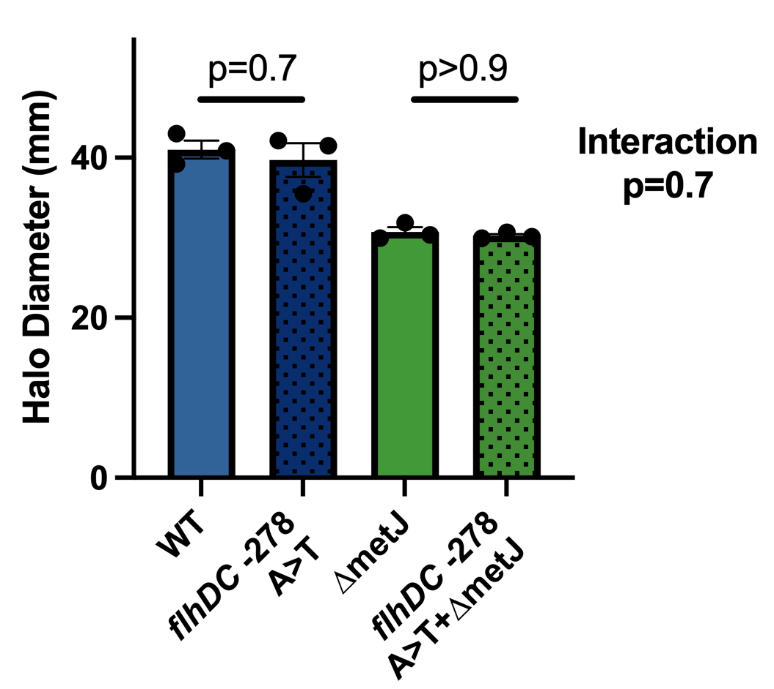

**Supplemental Figure 9: Methylation upstream of *flhDC* does not contribute to the ∆*metJ* motility defect.** The -278 GATC sequence is not required for the impacts of ∆*metJ* on motility. Motility on soft agar was measured six hours after inoculating the agar and following migration at 37°C, each dot represents the average of 3-5 technical replicates, data were normalized to the grand mean prior to plotting or performing statistics, and p-values were generated by two-way ANOVA with Sidak’s multiple comparisons test.

| **Supplemental Table 1: Bacterial strains used in this study** | | | |
| --- | --- | --- | --- |
| Identifier | Genotype | Plasmid | Antibiotic Resistance |
| DCK543 | *S.* Typhimurium NTCC 12023 (ATCC 14028s) |  |  |
| DCK545 | ∆*metJ* |  |  |
| DCK546 | *S.* Typhimurium NTCC 12023 (ATCC 14028s) | pWSK29 | Ampicillin |
| DCK547 | ∆*metJ* | pWSK29 | Ampicillin |
| DCK571 | *S.* Typhimurium NTCC 12023 (ATCC 14028s) | p67GFP3.1 | Ampicillin |
| DCK573 | ∆*metJ* | p67GFP3.1 | Ampicillin |
| DCK574 | *S.* Typhimurium NTCC 12023 (ATCC 14028s) | pWSK129 | Kanamycin |
| DCK576 | ∆*metJ* | pWSK129 | Kanamycin |
| DCK707 | ∆*dam* |  |  |
| DCK708 | ∆*dam*∆*metJ* |  |  |
| DCK711 | ∆*dam* | p67GFP3.1 | Ampicillin |
| DCK712 | ∆*dam*∆*metJ* | p67GFP3.1 | Ampicillin |
| DCK857 | ∆*dam* | pWSK29 | Ampicillin |
| DCK858 | ∆*dam*∆*metJ* | pWSK29 | Ampicillin |
| DCK859 | ∆*dam* | pWSK129 | Kanamycin |
| DCK860 | ∆*dam*∆*metJ* | pWSK129 | Kanamycin |
| DCK874 | ∆*dam* | pWSK129::*dam* | Kanamycin |
| DCK875 | ∆*dam*∆*metJ* | pWSK129::*dam* | Kanamycin |
| DCK934 | ∆*yhdJ* |  |  |
| DCK935 | ∆*metJ*∆*yhdJ* |  |  |
| DCK936 | ∆*yhdJ* | p67GFP3.1 | Ampicillin |
| DCK937 | ∆*metJ*∆*yhdJ* | p67GFP3.1 | Ampicillin |
| DCK938 | ∆*yhdJ* | pWSK29 | Ampicillin |
| DCK1015 | ∆*metJ*∆*yhdJ* | pWSK29 | Ampicillin |
| DCK939 | ∆*metJ*∆*yhdJ* | pWSK129 | Kanamycin |
| DCK865 | *flhC::3xFLAG* |  |  |
| DCK866 | *∆metJ*; *flhC::3xFLAG* |  |  |
| DCK1152 | *flhDC* promoter -278 A>T |  |  |
| DCK1155 | ∆*metJ*; *flhDC* promoter -278 A>T |  |  |
| DCK1157 | *∆tsr::Kan^R^* |  | Kanamycin |
| DCK1158 | ∆*metJ*∆*tsr::Kan^R^* |  | Kanamycin |

| **Supplemental Table 2: Plasmids used in this study** | | | |
| --- | --- | --- | --- |
| Identifier | Plasmid | Antibiotic Resistance | Source |
| DCK482 | pWSK29 | Ampicillin | (5) |
| DCK827 | pWSK129 | Kanamycin | (5) |
| DCK22 | P67GFP3.1 | Ampicillin | (6) |
| CS946 | pKD4 | Kanamycin | (7) |
| DCK599 | pKD46 | Ampicillin | (7) |
| CS943 | pCP20 | Ampicillin | (7) |
| DCK494 | pSUB11 | Ampicillin, Kanamycin | (8) |
| DCK1094 | pREDTKI | Kanamycin | (9) |
| DCK1095 | pMDIAI | Ampicillin | (9) |
| DCK1096 | pKSI-1 | Ampicillin | (9) |
| DCK855 | pWSK129::*dam* (includes 180 bases upstream and 5 bases downstream) | Kanamycin | This work |

| **Supplemental Table 3: Oligonucleotides used in this study** | | | |
| --- | --- | --- | --- |
| Name | Sequence (5’ 🡪 3’) | Purpose | Source |
| *flhD* Forward | TGTTCCGCCTCGGTATCAAC | qPCR | (10) |
| *flhD* Reverse | CGCGAATCCTGAGTCAAACG | qPCR | (10) |
| *dam* KD4 Forward | CTTTCTCCACAGCCGGAGAAGGTGTAATTAGTTAGTCAGCGTGTAGGCTGGAGCTGCTTC | Kanamycin Cassette | This study |
| *dam* KD4 Reverse | GGGGCAATCAAATACTGTTTCATCCGCTTCTCCTTGAGAACATATGAATATCCTCCTTAG | Kanamycin Cassette | This study |
| *yhdJ* KD4 Forward | GGGAAGCGCC TTTTTTATACGCATCACATG GAATTTGGTCGTGTAGGCTGGAGCTGCTTC | Kanamycin Cassette | This study |
| *yhdJ* KD4 Reverse | ACATGAAATTTTGACGCTGAAAAGCGGACTTACAATGCTTCATATGAATATCCTCCTTAG | Kanamycin Cassette | This study |
| *tsr* KD4 Forward | AGGCCGAAAATTCTGTATCTGTCTAGCGGAAAGAGAAAACGTGTAGGCTGGAGCTGCTTC | Kanamycin Cassette | This study |
| *tsr* KD4 Reverse | CCTACGGTCGTCTGTAGGCCGACTGTTCACCACTACGCCCCATATGAATATCCTCCTTAG | Kanamycin Cassette | This study |
| *flhC* pSUB11 Forward | TATTCCACAACTGCTGGATGAACAGATCGAACAGGCTGTTGACTACAAAGACCATGACGG | 3xFLAG tag | This study |
| *flhC* pSUB11 Reverse | TGACTTACCGCTGCTGGAGTGTTTGTCCACACCGTTTCGGCATATGAATATCCTCCTTAG | 3x FLAG tag | This study |
| *flhD* pMDIAI Forward | GGTTATTAATTAAACAAAGTAAAAGCCATGCTGATGGGTTCCCGGCGATCCTCTGG | Apramycin Cassette | This study |
| *flhD* pMDIAI Reverse | GAGATTCGCCTTACACGTTTACATCAATTTTTACAAATGTTGCATGACGGCAAGTGGACG | Apramycin Cassette | This study |
| *flhDC* pKSI-1 Forward | GATCTATCGAGGATCCTTTAGCTTTACTCTGTTTATCGCATTTCTGC | Plasmid Generation | This study |
| *flhDC* pKSI-1 Reverse | ATCGTAGTCTGTCGACGACATCATCCTTCCGCTGTTGACTATGAC | Plasmid Generation | This study |
| *flhDC* -278 A>T | GATTTTAGAAAATATGTGATGCAGAACACATATTTTAACGGAATACTTACGATAA | Site Directed Mutagenesis | This study |
| *flhDC* -278 A>T | TTATCGTAAGTATTCCGTTAAAATATGTGTTCTGCATCACATATTTTCTAAAATC | Site Directed Mutagenesis | This study |
| *dam* pWSK129 Forward | GATCTATCGAGGATCCTCCAGGCTGTGTCCTGCAATTGCCTGTGAGTGTC | Plasmid Generation | This study |
| *dam* pWSK129 Reverse | CGTAGCGTAAAAGCTTGAGAATTATTTTCTTGCAGGCGTTGCGACTCC | Plasmid Generation | This study |

| **Supplemental Table 4: Percent methylation compared to previous hypomethylation studies** | | | | | | | | |
| --- | --- | --- | --- | --- | --- | --- | --- | --- |
| **^Reproduced and Adapted from Sánchez-Romero *et al*. (4)** | | | | | | | | |
| **Gene^** | **Gene Product^** | **Number of GATCs^** | **Number of under-methylated GATCs^** | **Position(s) of GATC undermethylated A(s)^** | **Average % Methylation in wild-type bacteria grown in LB in this study (Combined Dataset)**  ***N/A = Could not find in dataset** | **Average % Methylation in ∆*metJ* bacteria grown in LB in this study (Combined Dataset)**  ***N/A = Could not find in dataset** | **% Methylation in wild-type SPI-2 media (Experiment 1)**  ***N/A = Count not find in dataset** | **% Methylation in ∆metJ SPI-2 media (Experiment 1)**  ***N/A = Count not find in dataset** |
| carA | Carbamoyl-phosphate synthase small chain | 2 | 2 | −511, −206 | **-511** = 52%  **-206** = 45.5% | **-511** = 49.5%  **-206** = 45% | **-511** = 49%  **-206** = 88% | **-511** = 69%  **-206** = 99% |
| dgoR | Galactonate operon transcriptional repressor | 1 | 1 | −147 | **-147** = 0% | **-147** =15.5% | **-147** = 63% | **-147** = 73% |
| ftnB | Ferritin-like protein | 4 | 2 | −340, −156 | **-340** = **N/A***  **-156** = 100% | **-340** = **N/A***  **-156** = 96.5% | **-340** = **N/A***  **-156** = 100% | **-340** = **N/A***  **-156** = 100% |
| gtr | O-antigen glycotransferase | 4 | 2 | −69, −56 | **N/A*** | **N/A*** | **N/A*** | **N/A*** |
| holA | DNA polymerase III, delta subunit | 5 | 1 | 4 | **+4** = 98.5% | **+4** = 97.5% | **+4** = 100% | **+4** = 100% |
| nanA | N-acetylneuraminate lyase | 8 | 1 | −58 | **-59*** = 13%  *No -58 GATC site in our mapping | **-59*** = 30.5%  *No -58 GATC site in our mapping | **-59*** = 0%  *No -58 GATC site in our mapping | **-59*** = 0%  *No -58 GATC site in our mapping |
| opvAB | O-antigen chain length regulation | 4 | 2 | −178, −105 | **N/A*** | **N/A*** | **N/A*** | **N/A*** |
| slrA | Glucitol/sorbitol-specific enzyme IIC component | 2 | 1 | −86 | **N/A*** | **N/A*** | **N/A*** | **N/A*** |
| ssaN | Type III secretion ATP synthase | 2 | 1 | −202 | **-202** = 100% | **-202** = 100% | **-202** = 100% | **-202** = 100% |
| STM1290 | N-acetylmannosamine-6-phosphate-2-epimerase | 2 | 1 | −465 | **N/A*** | **N/A*** | **N/A*** | **N/A*** |
| STM2047 | Hypothetical protein | 3 | 1 | −68 | **N/A*** | **N/A*** | **N/A*** | **N/A*** |
| STM3726 | Putative mannitol dehydrogenase | 3 | 1 | −68 | **N/A*** | **N/A*** | **N/A*** | **N/A*** |
| STM4889 | Putative Na^+^/galactosidase symporter | 4 | 1 | −171 | **N/A*** | **N/A*** | **N/A*** | **N/A*** |
| STM5047 | Putative cytoplasmic protein | 3 | 1 | −108 | **N/A*** | **N/A*** | **N/A*** | **N/A*** |
| STM5308 | Sugar transporter | 3 | 1 | −66 | **N/A*** | **N/A*** | **N/A*** | **N/A*** |
| yihU | Hypothetical oxidoreductase | 2 | 1 | −66 | **N/A*** | **N/A*** | **N/A*** | **N/A*** |
